# Deep learning enables therapeutic antibody optimization in mammalian cells by deciphering high-dimensional protein sequence space

**DOI:** 10.1101/617860

**Authors:** Derek M Mason, Simon Friedensohn, Cédric R Weber, Christian Jordi, Bastian Wagner, Simon Meng, Pablo Gainza, Bruno E Correia, Sai T Reddy

## Abstract

Therapeutic antibody optimization is time and resource intensive, largely because it requires low-throughput screening (10^3^ variants) of full-length IgG in mammalian cells, typically resulting in only a few optimized leads. Here, we use deep learning to interrogate and predict antigen-specificity from a massively diverse sequence space to identify globally optimized antibody variants. Using a mammalian display platform and the therapeutic antibody trastuzumab, rationally designed site-directed mutagenesis libraries are introduced by CRISPR/Cas9-mediated homology-directed repair (HDR). Screening and deep sequencing of relatively small libraries (10^4^) produced high quality data capable of training deep neural networks that accurately predict antigen-binding based on antibody sequence. Deep learning is then used to predict millions of antigen binders from an *in silico* library of ~10^8^ variants, where experimental testing of 30 randomly selected variants showed all 30 retained antigen specificity. The full set of *in silico* predicted binders is then subjected to multiple developability filters, resulting in thousands of highly-optimized lead candidates. With its scalability and capacity to interrogate high-dimensional protein sequence space, deep learning offers great potential for antibody engineering and optimization.

## INTRODUCTION

In antibody drug discovery, the ‘target-to-hit’ stage is a well-established process, as screening hybridomas, phage or yeast display libraries typically result in a number of potential lead candidates. However, the time and costs associated with lead candidate optimization often take up the majority of the preclinical discovery and development cycle^1^. This is largely due to the fact that lead optimization of antibody molecules consists of addressing multiple parameters in parallel, including expression level, viscosity, pharmacokinetics, solubility, and immunogenicity^2,3^. Once a lead candidate is discovered, additional engineering is often required; phage and yeast display offer a powerful method for high-throughput screening of large mutagenesis libraries (>10^9^), however they are primarily only used for increasing affinity or specificity to the target antigen^4^. The fact that nearly all therapeutic antibodies require expression in mammalian cells as full-length IgG means that the remaining development and optimization steps must occur in this context. Since mammalian cells lack the capability to stably replicate plasmids, this last stage of development is done at very low-throughput, as elaborate cloning, transfection and purification strategies must be implemented to screen libraries in the max range of 10^3^, meaning only minor changes (e.g., point mutations) are screened^5^. Interrogating such a small fraction of protein sequence space also implies that addressing one development issue will frequently cause rise of another or even diminish antigen binding altogether, making multi-parameter optimization very challenging. This challenge frequently results in antibodies with suboptimal biophysical properties for clinical development, which can lead to adverse side effects or even drug failure. For example, self-administered, subcutaneous injection of antibodies is becoming an increasingly used approach for patients requiring frequent dosing, but the identification of highly soluble, non-viscous antibodies which retain high biological activity is immensely difficult^6^. The withdrawal of Pfizer’s anti-PCSK9 antibody, bococizumab, from clinical trials is an even more drastic example, where the immunogenicity of the molecule adversely effected long-term treatment efficacy. Conversely, Sanofi and Regeneron’s clinically approved antibody, alirocumab, has the same molecular target of PCSK9, but shows almost no immunogenic effects^7^.

Machine learning applied to biological sequence data offers a powerful approach to augment protein engineering by constructing models capable of making predictions of genotype-phenotype relationships^8,9^. This is due to the capability of models to extrapolate complex relationships between sequence and function. One of the principle challenges in constructing accurate machine learning models is the collection of appropriate high-quality training data. Directed evolution platforms are well-suited for this as they rely on the linking of biological sequence data (DNA, RNA, protein) to a phenotypic output^10^. In fact, it has long been proposed to use machine learning models trained on data generated by mutagenesis libraries as a means to guide protein engineering^11,12^. Recently, Gaussian processes, a Bayesian learning model, were used to engineer cytochrome enzymes, enabling navigation through a vast protein sequence space to discover highly thermostable variants^13^. Similarly, the design and screening of a structure-guided library of channel rhodopsin membrane proteins was used to train Gaussian process regression models, which were able to accurately predict variants that could express and localize on mammalian cell membranes^14^.

In recent years, access to deep sequencing and parallel computing has enabled the construction of deep learning models capable of predicting molecular phenotype from sequence data^15,16^. For example, deep learning has been used to learn the sequence specificities of RNA- and DNA-binding proteins^17^, regulatory grammar of protein expression in yeast^18^, and HLA-neoantigen presentation on tumor cells^19^. In most cases deep (artificial) neural networks represent the class of algorithm utilized. While the complexity of neural networks has changed drastically since their conception, the fundamental concept remains the same: mimicking the connections of biological neurons to learn complex relationships between variables^20^. As an extension of a single-layer neural network, or perceptron^21^, deep learning incorporates multiple hidden layers to decipher relationships buried in large, high-dimensional data sets, such as the millions of reads gathered from a single deep sequencing experiment. Well trained models can then be used to make predictions on completely unseen and novel variants. This application of model extrapolation lends itself perfectly to protein engineering because it provides a way to interrogate a much larger sequence space than what is physically possible. For example, even for a short stretch of just 10 amino acids, the combinatorial sequence diversity explodes to 10^13^, a size which is nearly impossible to interrogate experimentally.

Here, we leverage the power of deep learning to perform multi-parameter optimization of therapeutic antibodies (full-length IgG) directly in mammalian cells (Figure 1). Starting with a mammalian display cell line^22^ expressing the therapeutic antibody trastuzumab (Herceptin), we use CRISPR-Cas9-mediated homology-directed repair (HDR) to introduce site-directed mutagenesis libraries in the variable heavy chain complementarity determining region 3 (CDRH3)^23^. In order to generate information rich training data, single-site deep mutational scanning (DMS) is first performed^24^, which is then used to guide the design of combinatorial mutagenesis libraries. An experimental (physical) library size of 5 × 10^4^ variants was then screened for specificity to the antigen HER2. All binding and non-binding variant sequences were then used to train recurrent and convolutional deep neural networks, which when fully-trained and optimized were able with high accuracy and precision to predict antigen-specificity based on antibody sequence. Neural networks are then used to predict antigen-specificity on a subset of sequence variants from the DMS-based combinatorial mutagenesis library (~10^8^ sequences), resulting in >3.0 × 10^6^ variants predicted to have a high probability of being antigen-specific. A random selection of variants were recombinantly expressed and tested, resulting in 30 out of 30 showing antigen-specific binding. The *in silico* library of predicted binders are then subjected to several sequence-based *in silico* filtering steps to optimize for developability parameters such as viscosity, clearance, solubility and immunogenicity, resulting in nearly 5,000 antibody sequence variants predicted to have more optimal properties than the starting trastuzumab sequence.

**Figure 1:**
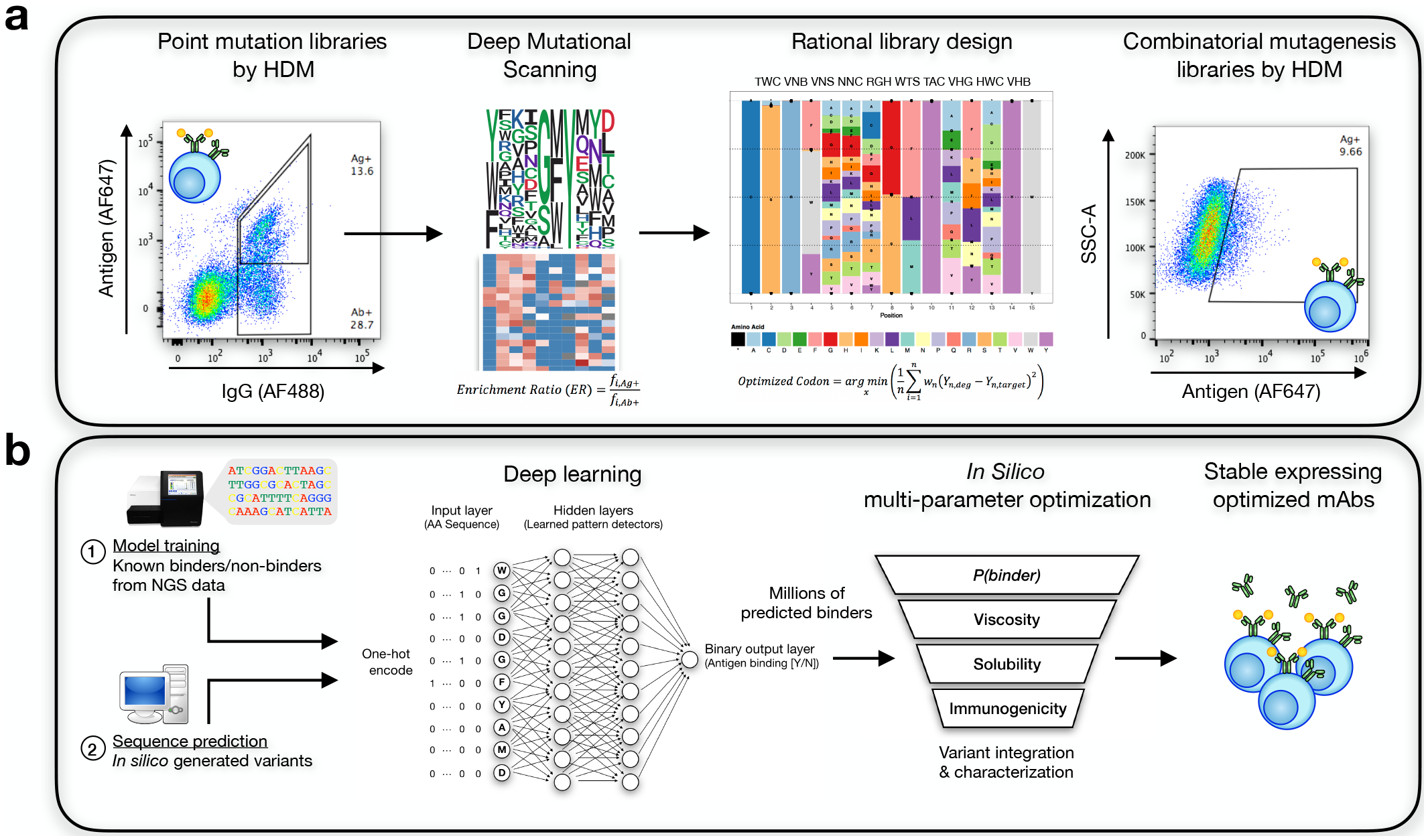
Implementing deep learning to predict antibody target specificity. **(a)** Generating quality data capable of training accurate models. First, deep mutational scanning assesses the impact mutations have on protein function across many different positions. These insights can then be applied to combinatorial mutagenesis strategies to guide protein library design capable of producing thousands of binding variants. **(b)** Sequence information for binders and non-binders can then be used to train deep neural networks to accurately predict antigen specificity of unknown antibody variants, producing millions of predicted binders. These binders can then be subjected to any available in silico methods for predicted various developability attributes.

## RESULTS

### Deep mutational scanning determines antigen-specific sequence landscapes and guides rational antibody library design

As the amino acid sequence of an antibody’s CDRH3 is a key determinant of antigen specificity, we performed DMS on this region to resolve the specificity determining residues. To start, a hybridoma cell-line was used that expressed a trastuzumab variant that could not bind HER2 antigen (mutated CDRH3 sequence) (Supplementary Fig. 1). Libraries were generated by CRISPR-Cas9-mediated homology-directed mutagenesis (HDM)^23^ by transfecting guide RNA (gRNA) targeting the CDRH3 and a pool of homology templates in the form of single-stranded oligonucleotides (ssODNs) containing NNK degenerate codons at single-sites tiled across CDRH3 (Figure 2a, Supplementary Fig. 2). Libraries were then screened by fluorescence activated cell sorting (FACS), and populations expressing surface IgG which either were binding or not binding to antigen were isolated and subjected to deep sequencing (Illumina MiSeq) (Supplementary Table 1). Deep sequencing data was then used to calculate enrichment scores of the 10 positions investigated, which revealed six positions that were sufficiently amenable to a wide-range of mutations and an additional three positions that were marginally accepting to defined mutations (Figure 2b). Although residues 102D, 103G, 104F, and 105Y appear to be contacting amino acids of the CDRH3 loop with HER2^25,26^, 105Y is the only residue completely fixed. In addition to DMS, we also explored the capacity of structural modeling to identify the prospective antigen-binding landscape by using structure-guided modeling with Rosetta, a leading software platform for computational protein design^27^. After performing 5,000 redesigns of trastuzamab (PDB id: 1N8Z), preceded by a stochastic backbone flexibility (FastRelax) step, 48 possible variants were generated (Figure 2c). The resulting sequence logo plot from these generated variants, however, differed substantially from the DMS-based sequence logo plot.

**Figure 2:**
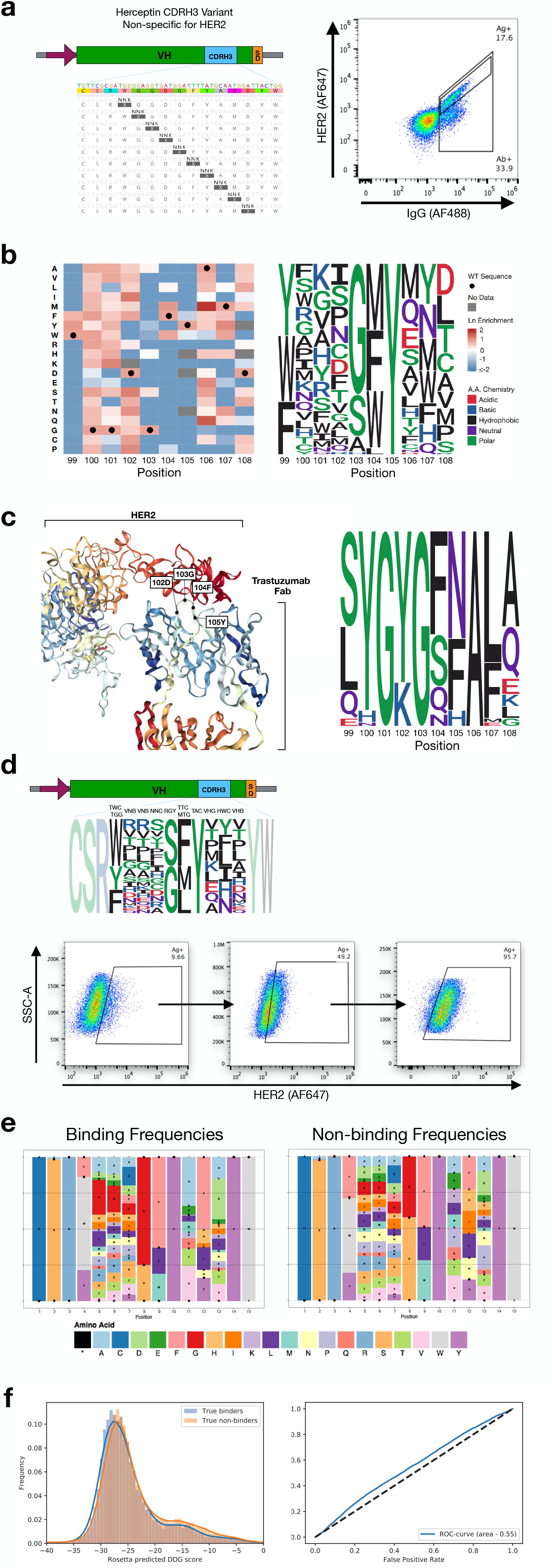
Sequence and structure-based analysis of the mutational landscape. **(a)** Flow cytometry profile following integration of tiled mutations by homology-directed mutagenesis. Antigen specific variants underwent 3 rounds of enrichment (Supplementary Fig. 2) **(b)** Corresponding heatmap (left) following sequencing analysis of the pre-sorted (Ab+) and post-sorted (Ag+) populations (Supplementary Table 1). Wild type amino acids are marked by black circles. The resulting sequence logo plot (right) generated by positively enriched mutations per position. **(c)** 3D protein structure of trastuzumab in complex with its target antigen, HER2^25,26^ (left). Locations of surface exposed residues: 102D, 103G, 104F, and 105Y are given. The protein design program Rosetta was run 5,000 times to generate sequence variants of trastuzumab predicted to bind the antigen HER2. The resulting sequence logo plot of the 48 generated CDRH3 loops (right) differs substantially from the DMS-based sequence logo plot. **(d)** Combinatorial mutagenesis libraries are designed from enrichment ratios observed in DMS data and integrated into the trastuzumab variant by homology-directed mutagenesis. Flow cytometry plots resulting from transfection of the rationally designed library. Deep sequencing was performed on the library (Ab+), non-binding variants (Ag−), and binding variants after 1 and 2 rounds of enrichment (Ag+1, Ag+2) (Supplementary Fig. 3, Supplementary Table 2). **(e)** Amino acid frequency plots of antigen binding and non-binding variants reveals nearly indistinguishable amino acid usages across all positions. **(f)** Distribution plot for the predicted Rosetta ddG scores for each sequence in the experimentally determined binding population (11,300 binders, shown in blue) and non-binding population (27,539 non-binders, shown in red) (left). The distribution plot is normalized for comparison purposes. Receiver operating characteristic (ROC) curve with area under curve (AUC) using the Rosetta ddG score to predict binding and non-binding variants shows poor classification (right).

The limited number of predictions and diversity from Rosetta suggested that sequence logo plots generated by DMS are a better option to guide the rational design of a combinatorial mutagenesis library, which consisted of degenerate codons across all positions (except 105Y) (Supplementary Fig. 3, Supplementary Table 7). Degenerate codons were selected per position based on their amino acid frequencies which most closely resembled the degree of enrichment found in the DMS data following 1, 2, and 3 rounds of antigen-specific enrichment (Supplementary Fig. 2, Equation 2). This combinatorial library possesses a theoretical protein sequence space of 7.17 × 10^8^, far greater than the single-site DMS library diversity of 200. Libraries containing CDRH3 variants were again generated in hybridoma cells through CRISPR-Cas9-mediated HDM in the same non-binding trastuzumab clone described previously (Figure 2d). Antigen binding cells were isolated by two rounds of enrichment by FACS (Figure 2d, Supplementary Fig. 3) and the binding/non-binding populations were subjected to deep sequencing. Sequencing data identified 11,300 and 27,539 unique binders and non-binders, respectively (Supplementary Table 2). These sequence variants represented only a miniscule 0.0054% of the theoretical protein sequence space of the combinatorial mutagenesis library.

Discriminating between the binding and non-binding sequences in the combinatorial library is challenging at the sequence level. Amino acid usage per position was comparatively similar between antigen binding and non-binding populations (Figure 2e), thus making it difficult to develop any sort of heuristic rules or decipher observable patterns to identify binding sequences. Thus, we investigated whether structure-based analysis could accurately predict antigen-binding sequences. We used Rosetta to model each of the 11,300 binder and 27,539 non-binder sequences from the combinatorial library on the antibody structure of trastuzumab, and used Rosetta’s predicted free energy of binding (ddG) as the discrimination score. This approach, however, yielded a very poor classifier (ROC curve AUC: 0.55, Figure 2f) and revealed that high-dimensional patterns determining antigen-specificity could not be extracted by structural modeling.

### Training deep neural networks to classify antigen-specificity based on antibody sequence

To learn the high-dimensional patterns that determine antigen binding, we set out to develop and train sequence-based deep learning models capable of predicting antibody specificity towards the target antigen HER2. After having compiled deep sequencing data on binding and non-binding CDRH3 variants, amino acid sequences were converted to an input matrix by one-hot encoding, an approach where each column of the matrix represents a specific residue and each row corresponds to the position in the sequence, thus a 10 amino acid CDRH3 sequence as here results in a 10 × 20 matrix. Each row will contain a single ‘1’ in the column corresponding to the residue at that position, whereby all other columns/rows receive a ‘0’. We utilized long short-term memory recurrent neural networks (LSTM-RNN) and convolutional neural networks (CNN), which represent two of the main classes of deep learning models used for biological sequence data^16^. LSTM-RNNs and CNNs both stem from standard neural networks, where information is passed along neurons that contain learnable weights and biases, however, there are fundamental differences in how the information is processed. LSTM-RNN layers contain loops, enabling information to be retained from one step to the next, allowing models to efficiently correlate a sequential order with a given output; CNNs, on the other hand, apply learnable filters to the input data, allowing it to efficiently recognize spatial dependencies associated with a given output. Model architecture and hyperparameters (Figures 3a, c) were selected by performing a grid search across various parameters (LSTM-RNN: nodes per layer, batch size, number epochs and optimizing function; CNN: number of filters, kernel size, dropout rate and dense layer nodes) using a k-fold cross-validation of the data set. All models were built to assess their accuracy and precision of classifying binders and non-binders from the available sequencing data. 70% of the original data set was used to train the models and the remaining 30% was split into two test data sets used for model evaluation: one test data set contained the same class split of sequences used to train the model and the other contained a class split of approximately 10/90 binders/non-binders to resemble physiological frequencies (Figure 2d). Performance of the LSTM-RNN and CNN were assessed by constructing receiver operating characteristic (ROC) curves and precision-recall (PR) curves derived from predictions on the unseen testing data sets (Figure 3b, d). Based on conventional approaches to training classification models, the data set was adjusted to allow for a 50/50 split of binders and non-binders during training. Under these training conditions, the LSTM-RNN and CNN were both able to accurately classify unseen test data (ROC curve AUC: 0.9 ± 0.0, average precision: 0.9 ± 0.0, Supplementary Fig. 5).

**Figure 3:**
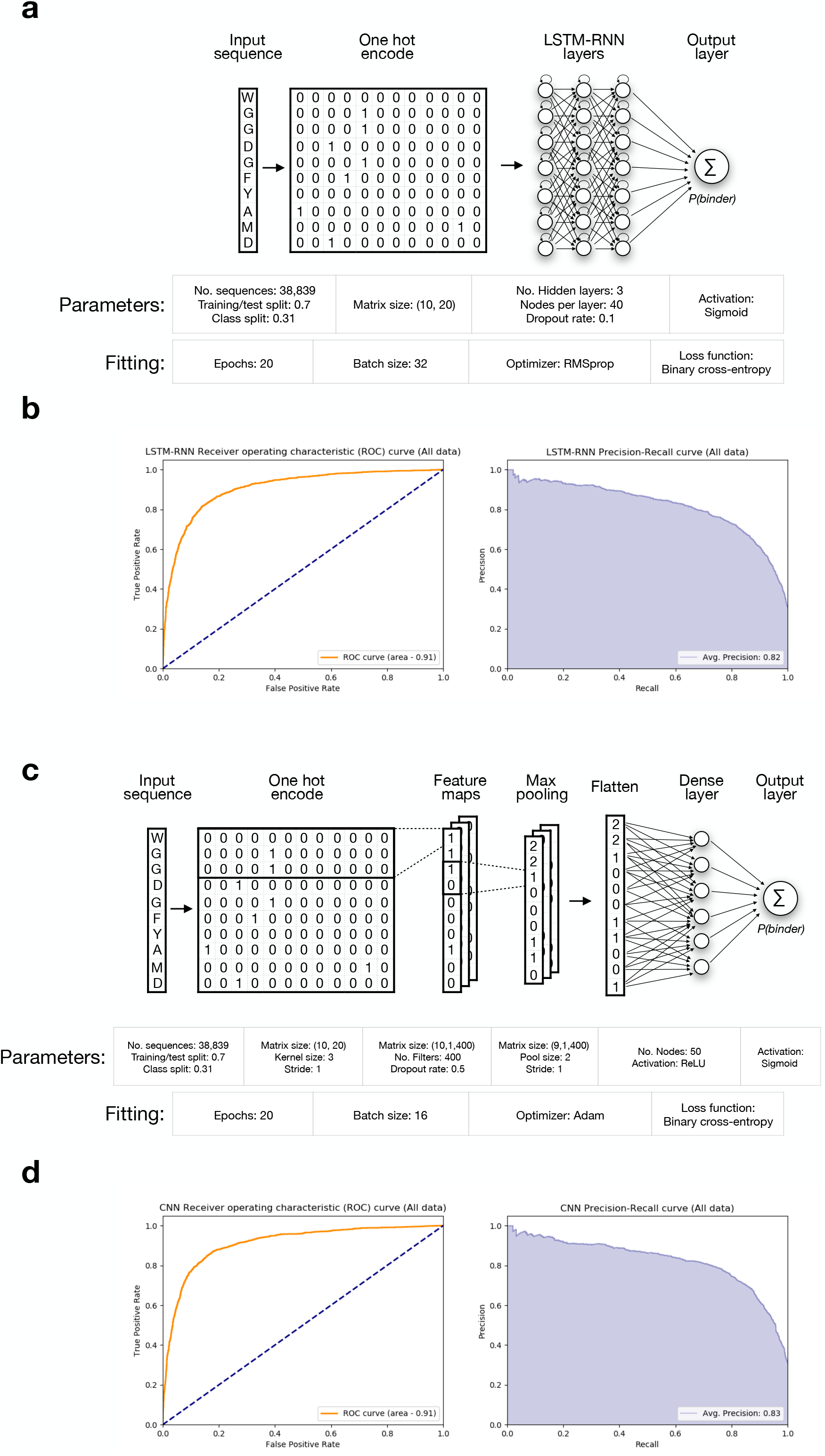
Deep learning models accurately predict antigen specificity. The selected network architectures and their model performance curves for classification of binding and non-binding sequences. Model training was performed on 70% of the data and testing was performed by withholding the remaining 30% and then comparing the model’s classification of test sequences with the known classification. In lieu of adjusting the data set to a defined class split of binding/ non-binding sequences, all known information was utilized to train and test the networks (approx. class split of 31%). **(a)** LSTM-RNN architecture and parameters used for model fitting. (**b**) ROC (receiver operating character) curve and PR (precision-recall) curve observed on the classification of sequences in the test set by the LSTM-RNN. (**c**) CNN architecture and parameters used for model fitting. (**d**) ROC curve and PR curve observed on the classification of sequences in the test set by the CNN. The high values observed for the ROC area under curve (AUC) and average precision of both networks represent robust measures of model accuracy and precision.

Next, we used the trained LSTM-RNN and CNN models to classify a random sample of 1 × 10^5^ sequences from the potential sequence space. We observed, however, an unexpectedly high occurrence of positive classifications (25,318 ± 1,643 sequences or 25.3 ± 1.6%, Supplementary Table 3b). With the knowledge that the physiological frequency of binders should be approximately 10-15%, we sought to adjust the classification split of the training data with the hypothesis that models were being subject to some unknown classification bias. Additional models were then trained on classification splits of both 20/80, and 10/90 binders/non-binders, as well as a classification split with all available data (approximately 30/70 binders/non-binders). Unbalancing the sequence classification led to a significant reduction in the percentage of sequences classified as binders, but also led to a reduction in the model performance on the unseen test data (Supplementary Fig. 4-7, Supplementary Tables 3a, b). Through our analysis, we concluded that the optimal data set for training the models was the set inclusive of all known CDRH3 sequences for the following reasons: 1) the percentage of sequences predicted as binders reflects this physiological frequency, 2) this data set maximizes the information the model sees, and 3) model performance on both test data sets. Final model architecture, parameters, and evaluation are shown in Figure 3. As a final measure of model validation, neural networks were trained with a data set containing randomly shuffled binding and non-binding class labels. Model performance of these networks revealed indiscriminate sequence classification on unseen test data (Supplementary Fig. 8), signifying the identification of learned patterns for networks trained with properly classified data.

### Predicted binding sequences are recombinantly expressed and antigen-specific

Using our DMS-based combinatorial mutagenesis library as a guide (Figure 2d), 7.2 ×10^7^ possible sequence variants were generated in silico. The fully-trained LSTM-RNN and CNN models were used to classify all 7.2 × 10^7^ sequence variants as either antigen binders or non-binders based on a probability score (*P*), resulting in a prediction of 8.55 × 10^6^ (LSTM-RNN) and 9.52 × 10^6^ (CNN) potential binders (*P > 0.50)*. This represented a reasonable fraction (11-13%) of antigen-specific variants based on experimental screening (Figure 2d). To increase confidence, we increased the prediction threshold for binder classification to *P* > 0.75 and took the consensus binders between the LSTM-RNN and CNN. This reduced the antigen-specific sequence space down to 3.1 × 10^6^ variants. To validate the precision of our fully trained LSTM-RNN and CNN models, we randomly selected and tested a subset of 30 CDRH3 sequences predicted to be antigen-specific (Figure 4a). To further demonstrate the capacity of deep learning to identify novel sequence variants, we also added the criteria that the selected variants must have a minimum Levenshtein distance (*LD)* of 5 from the original CDRH3 sequence of trastuzumab. CRISPR-Cas9-mediated HDR was used to generate mammalian display cell lines expressing the 30 different sequence variants. Flow cytometry was performed and revealed that 30 of the 30 variants (100%) were antigen-specific (Supplementary Fig. 9). Further analysis was performed on the 30 antigen-binding variants to more precisely quantify the binding kinetics via biolayer interferometry (BLI, FortéBio Octet RED96e) (Figure 4b). The original trastuzumab sequence was measured to have an affinity towards HER2 of 4.0 × 10^−10^ M (equilibrium dissociation constant, K_D_); and although the majority of variants tested had a slight decrease in affinity, 80% (24/30) were still in the single-digit nanomolar range, 17% (5/30) remained sub-nanomolar, and even one variant (3%) showed a near 3-fold increase in affinity compared to trastuzumab (K_D_ = 1.4 × 10^−10^ M) (Figure 4a, c). We also investigated if there were correlations between model prediction values and measured affinities (Supplementary Fig. 10). While no strong trend was observable, the highest affinity variants tended to have higher prediction values.

**Figure 4:**
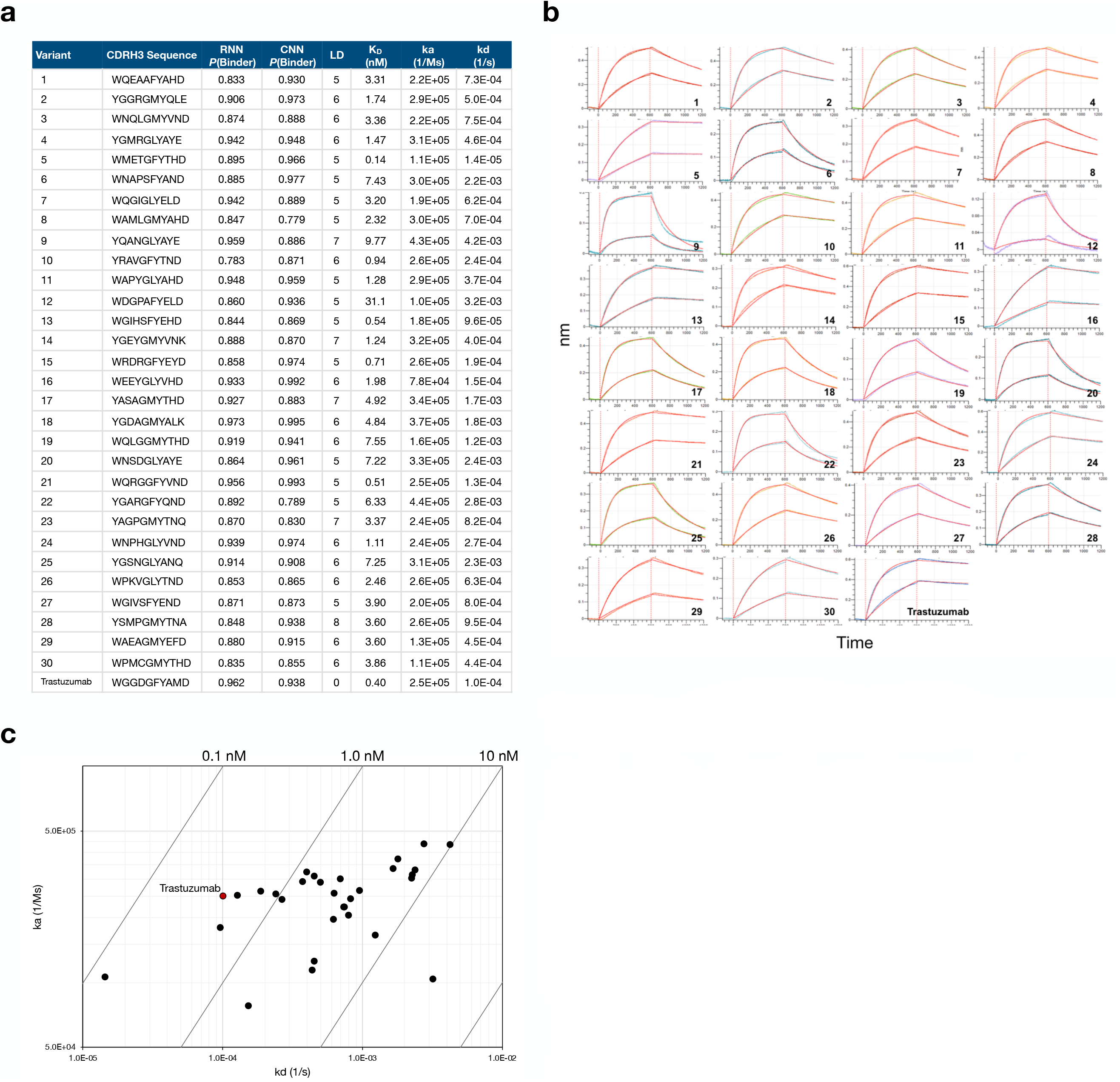
Deep neural network predicted sequences are experimentally validated to be antigen-specific. **(a)** To test the precision of the neural network predictions, 30 variants were randomly selected after increasing the prediction threshold (*P*(binder) > 0.75) and taking the consensus sequences between the LSTM-RNN and CNN. These sequences were integrated into individual hybridoma cells lines by separately transfecting ssODN donor sequences with gRNA. **(b)** Affinities for the 30 variant sequences were determined by biolayer interferometry (BLI). Although most sequences display a minor decrease in affinity for the target antigen, the majority of sequences still exude affinities of therapeutic relevance in the single nanomolar (24/30) or sub-nanomolar range (5/30). **(c)** Iso-affinity graph of the variant sequences.

### Sequence space analysis of deep learning predicted variants

In order to investigate the sequence space of the predicted binding and non-binding variants, we conducted a sequence similarity network analysis^28^ of 5,000 randomly selected binding and non-binding sequences (Supplementary Table 4, Supplementary Fig. 11). When generating similarity networks by clustering CDRH3 sequences with a *LD* ≤ 3, we observed 99.7% of all sequences to be within a single cluster, but when increasing the clustering stringency to a *LD* ≤ 2, the fraction of sequences found within the largest cluster is reduced to 30%, with the majority of other sequences not clustering with any other sequence (Figure 5a). While a large portion of the sequences found within the largest cluster are predicted binding sequences, non-binding sequences are also present, illustrating the complexities of the patterns identified by deep neural networks. To further elucidate the high-dimensional patterns of the antigen-binding landscape that deep learning models have identified, we performed the attribution method of *Integrated Gradients*^29^ on closely related sequences (*LD* ≤ 2) (Figure 5b). This analysis provides a means to visualize non-linear combinations of amino acids that contribute to classification as a binder or non-binder. This revealed that unlike position-weight matrices, LSTM-RNN and CNN models did not equally weight individual residues and positions and thus learned complex non-linear patterns associated with binding and non-binding.

**Figure 5:**
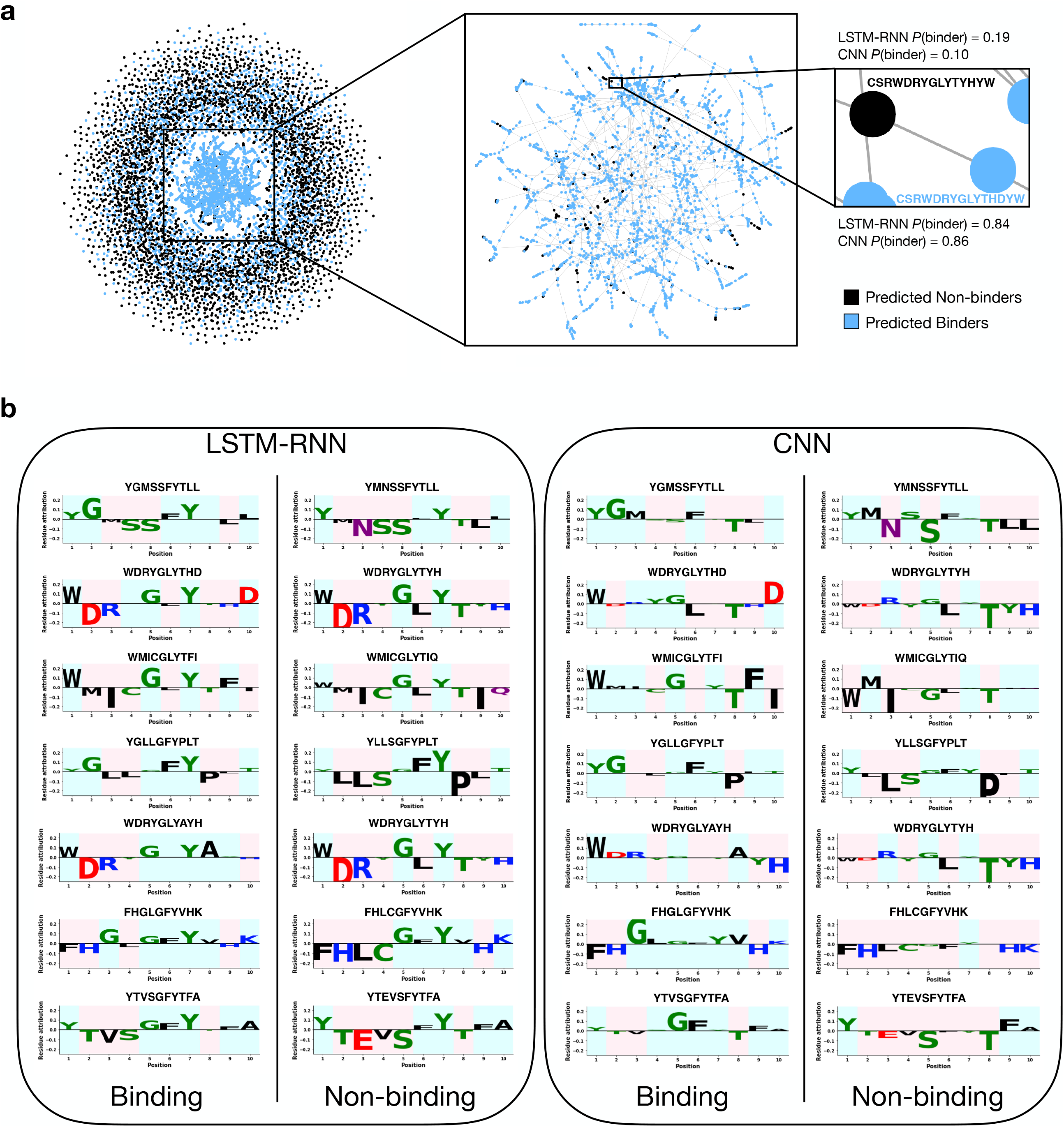
Deep neural networks decipher non-linear interactions to accurately classify binding and non-binding sequences. **(a)** A sequence similarity network analysis was completed on 5,000 randomly selected predicted binding (blue) and 5,000 non-binding variants (black) to investigate potential sequence similarities of the classification choices (left). Clustering was performed at a Levenshtein distance (LD) ≤ 2; Similarity network analyses performed with additional *LD*s can be found in Supplementary Table 4 and Supplementary Fig. 12. Although the largest cluster within the network (middle) contains 90% predicted binding variants, this comprises only 30% of all sequences in the network. Conversely, 42% of sequences do not cluster with any other neighboring sequences, thereby revealing that for the majority of variants, there are no discernible clusters of binding or non-binding predictions. **(b)** The Integrated Gradients method efficiently extracts and enables visualization of the classification patterns established by the LSTM-RNN (left) and the CNN (right). For the specific example, variants identified in the network with a *LD* of only 2 were classified as binding and non-binding sequences respectively. The LSTM-RNN and CNN uniquely identify non-linear combinations of amino acids that contribute to its classification as a binder (highlighted in green) or its classification as a non-binder (highlighted in red).

### Multi-parameter optimization for developability by in silico screening of antibody sequence space

Next, we characterized the full 3.1 × 10^6^ deep learning predicted antigen-specific sequences on a number of parameters to identify highly developable candidates compared to the original trastuzumab sequence. As a preliminary metric, we investigated their sequence similarity to the original trastuzumab sequence by calculating the *LD*. The majority of sequences showed an edit distance of *LD* > 4 (Figure 6a). The first step in filtering was to calculate the net charge and hydrophobicity index in order to estimate the molecule’s viscosity and clearance^2^. According to Sharma et al., viscosity decreases with increasing variable fragment (Fv) net charge and increasing Fv charge symmetry parameter (FvCSP); however, the optimal Fv net charge in terms of drug clearance is between 0 and 6.2 with a CDRL1+CDRL3+CDRH3 hydrophobicity index sum (HI sum) < 4. Based on the wide range of values for these parameters in the 3.1 × 10^6^ predicted variants (Figure 6b, c), we filtered any sequences out that had a FvCSP < 6.61 (trastuzumab FvCSP) or if they contained a Fv net charge > 6.2, and an HI sum > 4, < 0. This filtering criteria greatly reduced the sequence space down to 4.02 × 10^5^ variants. We next padded the CDRH3 sequences with 10 amino acids on the 5’ and 3’ ends and then ran these sequences through CamSol, a protein solubility predictor developed by Sormanni et al.^30^, which estimates and ranks sequence variants based on their theoretical solubility. The remaining variants produced a wide-range of protein solubility scores (Figure 6d) and sequences with a score < 0.5 (trastuzumab score) were filtered out, leaving 14,125 candidates for further analysis. As a last step in our *in silico* screening process, we aimed at reducing immunogenicity by predicting the peptide binding affinity of the variant sequences to MHC Class II molecules by utilizing NetMHCIIpan, a model previously developed by Jensen et al.^31^. One output from the model is a given peptide’s % Rank of predicted affinity compared to a set of 200,000 random natural peptides. Typically, molecules with a % Rank < 2 are considered strong binders and those with a % Rank < 10 are considered weak binders to the MHC Class II molecules scanned. All possible 15-mers from the padded CDRH3 sequences were run through NetMHCIIpan. After predicting the affinities for a set of 26 HLA alleles determined to cover over 98% of the global population^32^, sequences were filtered out if any of the 15-mers contained a % Rank < 5.5 (trastuzumab minimum % Rank) (Figure 6e). The number of 15-mers with a % Rank less than 10 and the average % Rank across all 15-mers for the remaining sequences were also calculated. Sequences with more than two 15-mers with a % Rank < 10 (Figure 6f) and those with an average % Rank < 60.56 (trastuzumab average % Rank) were also filtered out (Figure 6g). All remaining 4,881 variants contain values equal to or greater than the parameters of the original trastuzumab sequence. When applying this same filtering scheme on the 11,300 experimentally determined binding sequences (obtained from training / test data), only 9 variants remained. Lastly, to determine the best developable sequences, we calculated an overall developability improvement score based on the mean of normalized values for each relevant parameter (see Materials and Methods), where trastuzumab would have a developability improvement score equal to 0. Of the remaining 4,881 predicted binding sequences, 293 variants were identified to have a higher developability score compared to the maximum developability score of the 9 experimentally determined binding sequences (Figure 6h). The filtering parameters and number of remaining variants at each step for the *in silico* library are provided in Figure 6i.

**Figure 6:**
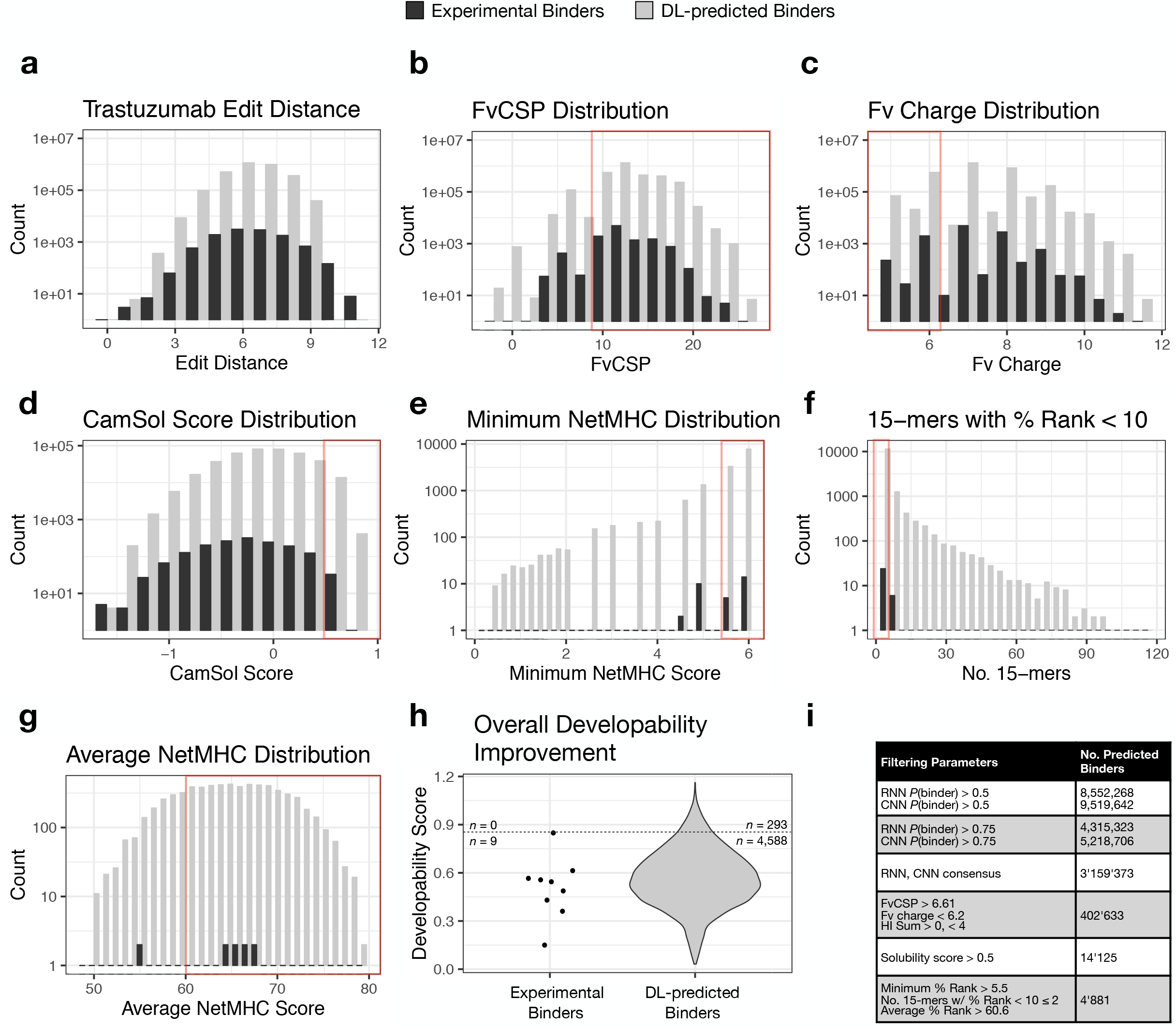
In silico screening of predicted binders identifies globally optimized variants. Antigen specific variants result in a wide range of *in silico* calculated parameters for developability. The following are staggered histograms showing the parameter distributions of all deep learning (DL)-predicted binders (light) and the experimentally observed binders (dark) at the different stages of filtering. Red boxes indicate filtering cut-offs determined by the developability metric calculated for the original trastuzumab sequence. **(a)** Levenshtein distance from wild-type trastuzumab. **(b)** Net charge of the VH domain. **(c)** CDRH3 hydrophobicity index. **(d)** CamSol intrinsic solubility score. **(e)** The minimum NetMHCIIpan % Rank (< 2 ~ strong affinity; < 10 ~ weak affinity) across all possible 15-mers for a given CDRH3 sequence and across all HLA alleles. **(f)** The number of 15-mers found within a given CDRH3 sequence that have a % Rank < 10 across all alleles. **(g)** The average NetMHCIIpan % Rank across all possible 15-mers and HLA alleles. **(h)** Scatter/violin plot for the overall developability improvement score (Eq. 3) of the remaining sequence variants passing all filtering criteria. 293 sequences of the predicted binders have a higher overall developability improvement score than the maximum score identified from an experimental binder. **(i)** Filtering parameters and the number of sequences at the corresponding stage of filtering.

## DISCUSSION

Addressing the limitation of antibody optimization in mammalian cells, we have developed an approach based on deep learning that enables us to identify antigen-specific sequences with high precision. Calculating and predicting various biophysical properties of antigen-specific variants allows for efficient identification of the most developable antibody molecule, resulting in significant time and cost savings and greatly reducing risk for downstream clinical development. Using the clinically approved antibody trastuzumab, we performed single-site DMS followed by combinatorial mutagenesis to determine the antigen-binding landscape of CDRH3. This DMS-based mutagenesis strategy is crucial for attaining high quality training data that is enriched with antigen-binding variants, in this case nearly 10% of our library (Figure 2d). In contrast, if a completely randomized combinatorial mutagenesis strategy was employed (i.e., NNK degenerate codons), it would be unlikely to produce any significant fraction of antigen-binding variants. In the future, other approaches to mutagenesis that generate enriched training data^33^, such as shotgun scanning mutagenesis^34^, binary substitution^35^ and recombination^14,36^ may also be explored for training deep neural networks.

Our initial single-site DMS libraries screened for enriched mutations through antigen-binding, yet combining these mutations in a cohesive manner to alter biophysical properties while retaining high antigen affinity is challenging. The amino acid composition of binding and non-binding variants is highly similar (Figure 2e), and visually identifying the sequence patterns that lead to binding is a daunting, if not impossible task. Moreover, structure-based modeling was unable to discriminate between binders and non-binders as predicting fine-grained protein-complex affinities is highly challenging using generalistic methods such as Rosetta^27^. This is compounded by introducing CDRH3 loop mutations which likely result in challenging loop conformational changes^37^. While more advanced, ensemble-based ddG prediction methods^38^ could result in better performance, applying this to millions of sequences may be infeasible, further exemplifying the value of deep neural networks that are able to learn the high-dimensional space of antigen-binding sequences.

A remarkable finding in this study was that experimental screening of a library of only 5 × 10^4^ variants, which reflected a tiny fraction (0.0054%) of the total sequence diversity of the DMS-based combinatorial mutagenesis library (7.17 × 10^8^), was capable of training accurate neural networks. This suggests that physical library size limitations of mammalian expression systems (or other expression platforms such as phage and yeast) and deep sequencing read depth will not serve as a limitation for deep learning-guided protein engineering. Another important result was that deep sequencing of antigen-binding and non-binding populations showed nearly no observable difference in their positional amino acid usage (Figure 2e), revealing that neural networks are effectively capturing high-dimensional and non-linear patterns/interactions (Figure 5b).

In the current study, we selected LSTM-RNNs and CNNs as the basis of our classification models, as they represent two state of the art approaches in deep learning. Other machine learning approaches such as k-nearest neighbors, random forests, and support vector machines are also well-suited at identifying complex patterns from input data, but as data set sizes continue to grow, as is realizable with biological sequence data, deep neural networks tend to outperform these classical techniques^17^. Furthermore, deep generative modeling methods such as variational autoencoders and generative adversarial networks may also be used to explore the mutagenesis sequence space from directed evolution^39^.

We *in silico* generated approximately 7.2 × 10^7^ CDRH3 variants from DMS-based combinatorial diversity and used fully trained LSTM-RNN and CNN models to classify each sequence as a binder or non-binder. The 7.2 × 10^7^ sequence variants comprise only a subset of the potential sequence space and was chosen to minimize the computational effort, however, it still represents a library size several orders of magnitude greater than what is experimentally achievable in mammalian cells. We easily envision extending the screening capacity through script optimization and employing parallel computing on high performance clusters. Out of all variants classified, the LSTM-RNN and CNN predicted approximately 11-13% to bind the target antigen, showing exceptional agreement with the experimentally observed frequencies by flow cytometry (Figure 2d). In order to experimentally validate the precision of neural networks to predict antigen specificity, we randomly selected and expressed 30 variants from the library of sequences with a minimum edit distance of 5 from trastuzumab. The precision of the LSTM-RNN and CNN models were each estimated to be ~85% (at *P* > 0.75) according to predictions made on the test data sets (Figure 3b, d). By taking the consensus between models, however, we experimentally validated that all randomly selected (30/30) of the antigen-predicted sequences were indeed binders, and several of which were high affinity. While we anticipate false positives would be observed by increasing the sample size tested, validation of this subset strongly infers that potentially thousands of optimized lead candidates maintain a binding affinity in the range of therapeutic relevance, while also containing substantial sequence variability from the starting trastuzumab sequence. Future work to increase the stringency of selection during screening or a more detailed investigation of correlations between prediction probability and affinity could prove insightful towards retaining high target affinities. Experimentally validating the accuracy of the models to predict the binding status of sequence variants led us to take a more in depth look at the sequence space of predicted binding and non-binding variants. A sequence similarity network analysis at various *LD*s revealed no distinct clusters between binding and non-binding sequences, indicating an overall sequence similarity of both classifications. By then quantitatively analyzing neural network predictions, we were able to shed light on the high-dimensional patterns captured by the respective models and decipher amino acid combinations contributing to a sequence’s classification.

Once an antibody’s affinity for its target antigen is within a desirable range for efficacious biological modification, addressing other biophysical properties becomes the focus of antibody development. With recent advances in computational predictions^40,41^, a number of these properties, including viscosity, clearance, stability^2^, specificity^42^, solubility^30^ and immunogenicity^31^ can be approximated from sequence information alone. With the aim of selecting antibodies with improved characteristics, we subjected the library of predicted binders to a number of these *in silico* approaches in order to provide a ranking structure and filtering strategy for developability (Figure 6). After implementing these methods to remove variants with a high likelihood of having poor viscosity, clearance or solubility, as well as those with high immunogenic potential, nearly 5,000 multi-parameter optimized antibody variants remained with developability scores greater than the original trastuzumab sequence. Although a limited number of developable sequences can be initially identified experimentally (Figure 6h), this only reflects a small fraction of the highly-developable sequence space (0.2%). By screening *in silico* libraries, the presence of every sequence variant within the defined space is guaranteed, ensuring the identification of globally optimized sequences. Future work to apply more stringent or additional filters which address other developability parameters (e.g. stability, specificity, humanization) could also be implemented to further reduce the sequence space down to the most developable therapeutic candidates across even more parameters. For instance, previous studies have investigated the likeness of therapeutic antibodies to the human antibody repertoire^43^. We also envision this approach to enable the optimization of other functional properties of therapeutic antibodies, such as pH-dependent antibody recycling^44^ or affinity/avidity tuning^45,46^. Additionally, extending this approach to other regions across the variable light and heavy chain genes, namely other CDRs, may yield deep neural networks that are able to capture long-range, complex relationships between an antibody and its target antigen. To explore these patterns in greater depth, it may be useful to compare neural network predictions with other advanced structural modeling techniques such as ones that take advantage of geometric deep learning^47^.

## METHODS

### Mammalian cell culture and transfection

Hybridoma cells were cultured and maintained according to the protocols described by Mason et al.^23^. Hybridoma cells were electroporated with the 4D-Nucleofector™System (Lonza) using the SF Cell Line 4D-Nucleofector^®^ × Kit L or × Kit S (Lonza, V4XC-2024, V4XC-2032) with the program CQ-104. Cells were prepared as follows: cells were isolated and centrifuged at 125 × G for 10 minutes, washed with Opti-MEM^®^ I Reduced Serum Medium (Thermo, 31985-062), and centrifuged again with the same parameters. The cells were resuspended in SF buffer (per kit manufacturer guidelines), after which Alt-R gRNA (IDT) and ssODN donor (IDT) were added. All experiments performed utilize constitutive expression of Cas9 from *Streptococcus pyogenes* (SpCas9). Transfections of 1×10^6^ and 1×10^7^ cells were performed in 100 μl, single Nucleocuvettes™ with 0.575 or 2.88 nmol Alt-R gRNA and 0.5 or 2.5 nmol ssODN donor respectively. Transfections of 2×10^5^ cells were performed in 16-well, 20 μl Nucleocuvette™ strips with 115 pmol Alt-R gRNA and 100 pmol ssODN donor.

### Flow cytometry analysis and sorting

Flow cytometry-based analysis and cell isolation were performed using the BD LSR Fortessa™ (BD Biosciences) and Sony SH800S (Sony), respectively. When labeling with fluorescently conjugated antigen or anti-IgG antibodies, cells were first washed with PBS, incubated with the labeling antibody and/or antigen for 30 minutes on ice, protected from light, washed again with PBS and then analyzed or sorted. The labeling reagents and working concentrations are described in Supplementary Table 5. For cell numbers different from 10^6^, the antibody/antigen amount and incubation volume were adjusted proportionally.

### Sample preparation for deep sequencing

Sample preparation for deep sequencing was performed similar to the antibody library generation protocol of the primer extension method described previously^48^. Genomic DNA was extracted from 1-5×10^6^ cells using the Purelink™ Genomic DNA Mini Kit (Thermo, K182001). Extracted genomic DNA was subjected to a first PCR step. Amplification was performed using a forward primer binding to the beginning of the VH framework region and a reverse primer specific to the intronic region immediately 3’ of the J segment. PCRs were performed with Q5® High-Fidelity DNA polymerase (NEB, M0491L) in parallel reaction volumes of 50 ml with the following cycle conditions: 98°C for 30 seconds; 16 cycles of 98°C for 10 sec, 70°C for 20 sec, 72°C for 30 sec; final extension 72°C for 1 min; 4°C storage. PCR products were concentrated using DNA Clean and Concentrator (Zymo, D4013) followed by 0.8X SPRIselect (Beckman Coulter, B22318) left-sided size selection. Total PCR1 product was amplified in a PCR2 step, which added extension-specific full-length Illumina adapter sequences to the amplicon library. Individual samples were Illumina-indexed by choosing from 20 different index reverse primers. Cycle conditions were as follows: 98°C for 30 sec; 2 cycles of 98°C for 10 sec, 40°C for 20 sec, 72°C for 1 min; 6 cycles of 98°C for 10 sec, 65°C for 20 sec, 72°C for 1 min; 72°C for 5 min; 4°C storage. PCR2 products were concentrated again with DNA Clean and Concentrator and run on a 1% agarose gel. Bands of appropriate size (~550bp) were gel-purified using the Zymoclean™ Gel DNA Recovery kit (Zymo, D4008). Concentration of purified libraries were determined by a Nanodrop 2000c spectrophotometer and pooled at concentrations aimed at optimal read return. The quality of the final sequencing pool was verified on a fragment analyzer (Advanced Analytical Technologies) using DNF-473 Standard Sensitivity NGS fragment analysis kit. All samples passing quality control were sequenced. Antibody library pools were sequenced on the Illumina MiSeq platform using the reagent kit v3 (2×300 cycles, paired-end) with 10% PhiX control library. Base call quality of all samples was in the range of a mean Phred score of 34.

### Bioinformatics analysis and graphics

The MiXCR v2.0.3 program was used to perform data pre-processing of raw FASTQ files^49^. Sequences were aligned to a custom germline gene reference database containing the known sequence information of the V- and J-gene regions for the variable heavy chain of the trastuzumab antibody gene. Clonotype formation by CDRH3 and error correction were performed as described by Bolotin et al^49^. Functional clonotypes were discarded if: 1) a duplicate CDRH3 amino acid sequence arising from MiXCR uncorrected PCR errors, or 2) a clone count equal to one. Downstream analysis was performed using R v3.2.2^50^ and Python v3.6.5^51^. Graphics were generated using the R packages ggplot2^52^, RColorBrewer^53^, and ggseqlogo^54^.

### Calculation of enrichment ratios (ERs) in DMS

The ERs of a given variant was calculated according to previous methods^55^. Clonal frequencies of variants enriched for antigen specificity by FACS, f_i,Ag+_, were divided by the clonal frequencies of the variants present in the original library, f_i,Ab+_, according to Equation 1.

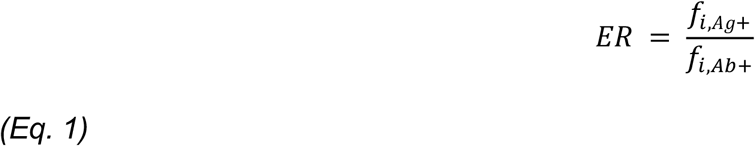

A minimum value of −2 was designated to variants with log[ER] values less than or equal −2 and variants not present in the dataset were disregarded in the calculation. A clone was defined based on the exact a.a. sequence of the CDRH3.

### Redesign of trastuzumab in Rosetta for diversity of sequences

The Rosetta program^27^ was used to redesign the trastuzumab antibody in complex with the extracellular domain of HER2 (PDB id: 1N8Z)^25^. Ten residues in the CDRH3 loop of trastuzumab (residues 98-108 of the heavy chain) were allowed to mutate to any natural amino acid, while all other residues were allowed to change rotameric conformation. A RosettaScript invoked the PackRotamersMover, a stochastic MonteCarlo algorithm, to optimize the sequence of the antibody to CDRH3 according to the Rosetta energy function, followed by backbone minimization. Energies were computed using Rosetta's ddG filter. Rosetta was run to generate 5000 sequences stochastically, and this resulted in 48 sequences. Rosetta's output files were processed using RS-Toolbox^56^.

### Classification of experimentally-determined sequences in Rosetta

Each of the 11,300 binding and 27,539 non-binding sequences from the combinatorial library were modelled in Rosetta^27^. For each experimentally-determined binding or non-binding sequence, the structure of the HER2:trastuzumab complex was used as input and the residues diverging from the wildtype were mutated using the PackRotamersMover in RosettaScripts^57^. The backbone and the side chains were minimized with Rosetta's MinMover after the sequence was modeled to optimized intra- and inter-chain contacts. Rosetta's predicted interface score (ddG) was used as the relative classification score.

### Codon selection for rational library design

Codon selection for rational library design was based off the equation provided by Mason et al.^23^, (Equation 2), where *Y*_*n,deg*_ represents the amino acid frequency for a given degenerate codon scheme, *Y*_*n,target*_ is the target amino acid frequency, and *n* is the number of amino acids, 20. Residues identified in DMS analysis to have a positive enrichment (ER > 1, or log[ER] > 0) were normalized according to their enrichment ratios and were converted to theoretical frequencies and taken as the target amino acid frequencies. Degenerate codon schemes were then selected which most closely reflect these frequencies as calculated by the mean squared error between the degenerate codon and the target frequencies.

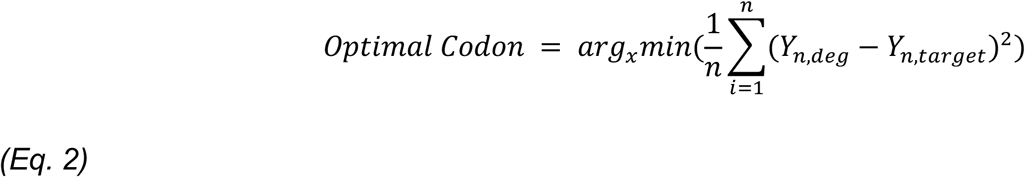

In certain instances, if the selected degenerate codon did not represent desirable amino acid frequencies or contained undesirable amino acids, a mixture of degenerate codons were selected and pooled together to achieve better coverage of the functional sequence space.

### Deep learning model construction

Machine learning models were built in Python v3.6.5. LSTM-RNNs, and CNNs were built using the Keras^58^ v2.1.6 Sequential model as a wrapper for TensorFlow^59^ v1.8.0. Model architecture and hyperparameters were optimized by performing a grid search of relevant variables for a given model. These variables include nodes per layer, activation function(s), optimizer, loss function, dropout rate, batch size, number of epochs, number of filters, kernel size, stride length, and pool size. Grid searches were performed by implementing a k-fold cross validation of the data set.

### Deep learning model training and testing

Data sets for antibody expressing, non-binding, and binding sequences (Sequencing statistics: Supplementary Tables 1, 2) were aggregated to form a single, binding/non-binding data set where antibody expressing sequences were classified as non-binders, unless also identified among the binding sequences. Sequences from one round of antigen enrichment were excluded from the training data set. The complete, aggregated data set was then randomly arranged and appropriate class labeled sequences were removed to achieve the desired classification ratio of binders to non-binders (50/50, 20/80, 10/90, and non-adjusted). The class adjusted data set was further split into a training set (70%), and two testing sets (15% each), where one test set reflected the classification ratio observed for training and the other reflected a classification ratio of approximately 10/90 to resemble the physiological expected frequency of binders.

### Sequence similarity and model attribution analysis of predicted variants

Sequence similarity networks of sequences predicted to be antigen positive and antigen negative were constructed for Levenshtein Distance 1-6 were constructed using the igraph R package^60^ v1.2.4. The resulting networks were analyzed with respect to their overall connectivity, the composition of their largest clusters and the overall degree distribution between the classes.

The Integrated Gradients technique^29^ was used to assess the relative attribution of each feature of a given input sequence towards the final prediction score. First, a baseline was obtained by zeroing out the input vector and the path integral of the gradients from baseline to the input vector was then approximated with a step size of 100. Integrated gradients were visualized as sequence logos. Sequence logos were created by the python module Logomaker^61^.

### In silico sequence classification and sequence parameters

All possible combinations of amino acids present in the DMS-based combinatorial mutagenesis libraries were used to calculate the total theoretical sequence space of 7.17 × 10^8^. 7.2 × 10^7^ sequence variants were generated *in silico* by taking all possible combinations of the amino acids used per position in the combinatorial mutagenesis library designed from the DMS data following three rounds of enrichment for antigen binding variants (Supplementary Fig. 2c, 3c); Alanine was also selected to be included at position 103. All *in silico* sequences were then classified as a binder or non-binder by the trained LSTM-RNN and CNN models. Sequences were selected for further analysis if they were classified in both models with a prediction probability (*P*) of more than 0.75.

The Fv net charge and Fv charge symmetry parameter (FvCSP) were calculated as described by Sharma et al. Briefly, the net charge was determined by first solving the Henderson-Hasselbalch equation for each residue at a specified pH (here 5.5) with known amino acid pKas^62^. The sum across all residues for both the VL and VH was then calculated as the Fv net charge. The FvCSP was calculated by taking the product of the VL and VH net charges. The hydrophobicity index (HI) was also calculated as described by Sharma et al., according to the following equation: HI = −(∑n_i_E_i_ / ∑n_j_E_j_). E represents the Eisenberg value of an amino acid, n is the number of an amino acid, and i and j are hydrophobic and hydrophilic residues respectively.

The protein solubility score was determined for each, full-length CDRH3 sequence (15 a.a.) padded with 10 amino acids on both the 5’ and 3’ ends (35 a.a.) by the CamSol method^30^ at pH 7.0.

The binding affinities for a reference set of 26 HLA alleles^32^ were determined for each 15-mer contained within the 10 amino acid padded CDRH3 sequence (35 a.a.) by NetMHCIIpan 3.2^31^. The output provides for each 15-mer a predicted affinity in nM and the % Rank which reflects the 15-mer’s affinity compared to a set of random natural peptides. The % Rank measure is unaffected by the bias of certain molecules against stronger or weaker affinities and is used to classify peptides as weak or strong binders towards the specified MHC Class II allele. The minimum % Rank, the number of 15-mers with % Rank less than 10 (classification of weak binder), and the average % Rank were calculated across all 21 15-mers for a single CDRH3 sequence across all 26 HLA alleles.

Overall developability improvement of the antibody sequences was determined by first normalizing the FvCSP, CamSol score, and average NetMHCII % Rank according to the range of values observed in the remaining sequences post-filtering. The normalized CamSol protein solubility score was then weighted by a factor of 2 for its importance in determining developability. Lastly, the mean across these three parameters was taken to produce the overall developability improvement score. Since the sequences were filtered with the calculated values for trastuzumab, trastuzumab would have an overall developability improvement equal to 0.

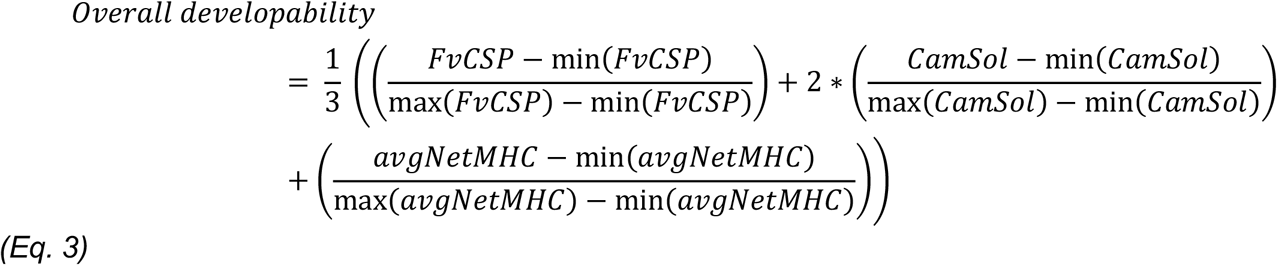

### Affinity measurements by biolayer interferometry

Monoclonal populations of the individual variants were isolated by performing a single-cell sort. Following expansion, supernatant for all variants was collected and filtered through a 0.20 μm filter (Sartorius, 16534-K). Affinity measurements were then performed on an Octet RED96e (FortéBio) with the following parameters. Anti-human capture sensors (FortéBio, 18-5060) were hydrated in conditioned media diluted 1 in 2 with kinetics buffer (FortéBio, 18-1105) for at least 10 minutes before conditioning through 4 cycles of regeneration consisting of 10 seconds incubation in 10 mM glycine, pH 1.52 and 10 seconds in kinetics buffer. Conditioned sensors were then loaded with 0 μg/mL (reference sensor), 10 μg/mL trastuzumab (reference sample), or hybridoma supernatant (approximately 20 μg/mL) diluted 1 in 2 with kinetics buffer followed by blocking with mouse IgG (Rockland, 010-0102) at 50 μg/mL in kinetics buffer. After blocking, loaded sensors were equilibrated in kinetics buffer and incubated with either 5 nM or 25 nM HER2 protein (Sigma-aldrich, SRP6405-50UG). Lastly, sensors were incubated kinetics buffer to allow antigen dissociation. Kinetics analysis was performed in analysis software Data Analysis HT v11.0.0.50.

## ACKNOWLEDGEMENTS

We acknowledge the ETH Zurich D-BSSE Single Cell Unit and the ETH Zurich D-BSSE Genomics Facility for support, in particular, M. Di Tacchio, A. Gumienny, E. Burcklen, and C. Beisel. We also thank the Vendruscolo Lab (Cambridge, UK), in particular P. Sormanni, for assistance with implementing the CamSol method on large libraries, as well as the group of Prof. Morten Nielson (DTU, Denmark) for providing an easy-to-use package for MHC Class II affinity predictions. Funding was provided by the National Competence Center for Research on Molecular Systems Engineering.

## AUTHOR CONTRIBUTIONS

D.M.M., S.F., C.R.W. and S.T.R. developed the methodology; D.M.M. and S.T.R. designed the experiments and wrote the manuscript; D.M.M., C.R.W. and S.F. analyzed sequencing data and performed deep learning analysis; P.G. and B.E.C. designed and performed structural modelling experiments and analysis; C.J. generated in silico libraries; D.M.M. performed experiments; B.W., and S.M.M. performed cell line development.

