## Supplementary Material for "Deep learning enables therapeutic antibody optimization in mammalian cells by deciphering high-dimensional protein sequence space" (from another version of this preprint: version 2, 2019-05-30)

### Supplementary Figures and Tables

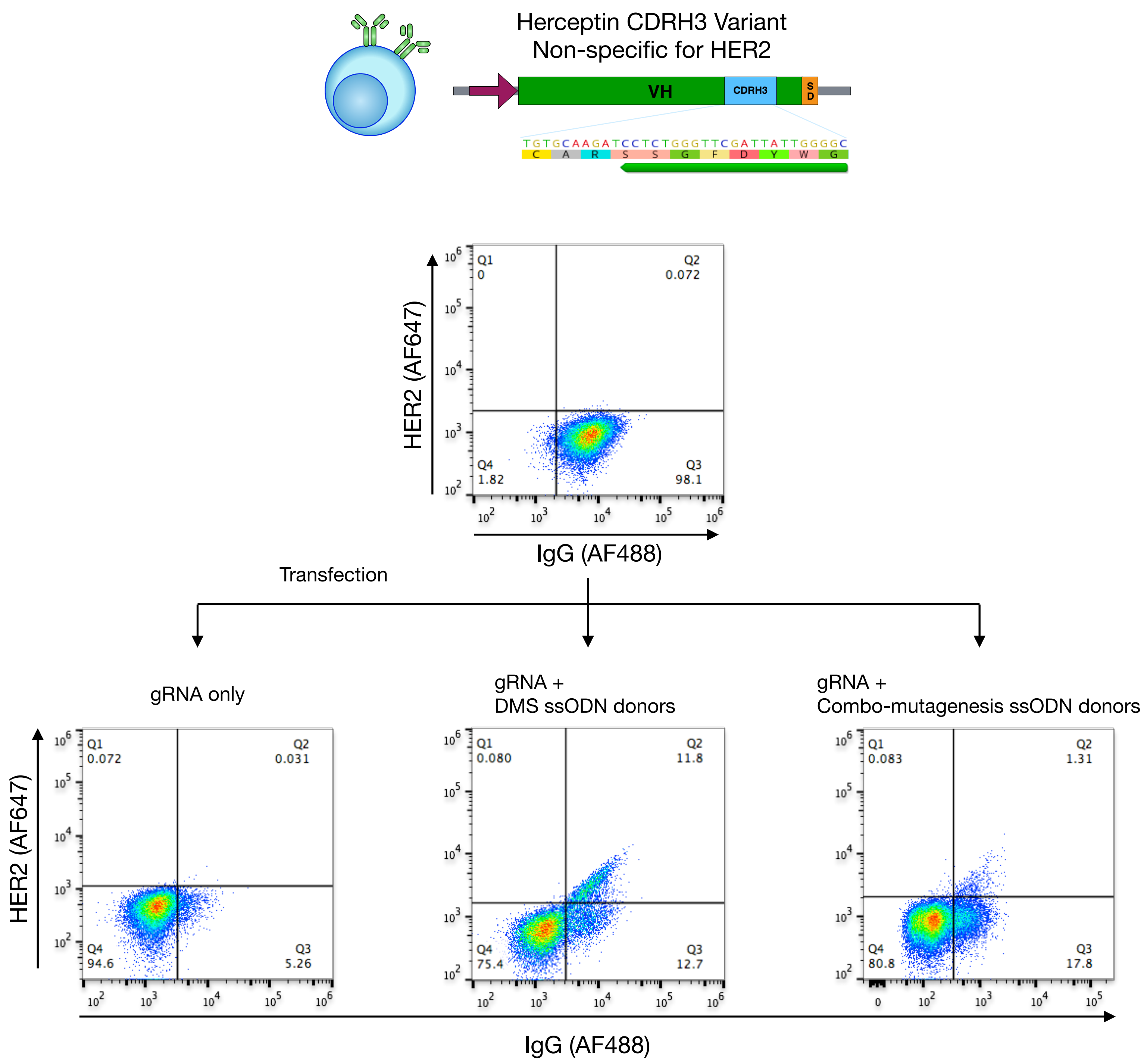


**Supplementary Figure 1: Trastuzumab CDRH3 (non-binding) variant**

A trastuzumab variant was generated by replacing the CDRH3 sequence with the sequence shown above, eliminating binding to the HER2 protein. Flow cytometry profile of the trastuzumab variant (top) and the resulting flow cytometry profiles following transfection of the trastuzumab variant with either gRNA only (left), gRNA+DMS ssODN library (middle), or gRNA+DMS-combinatorial mutagenesis library (right).

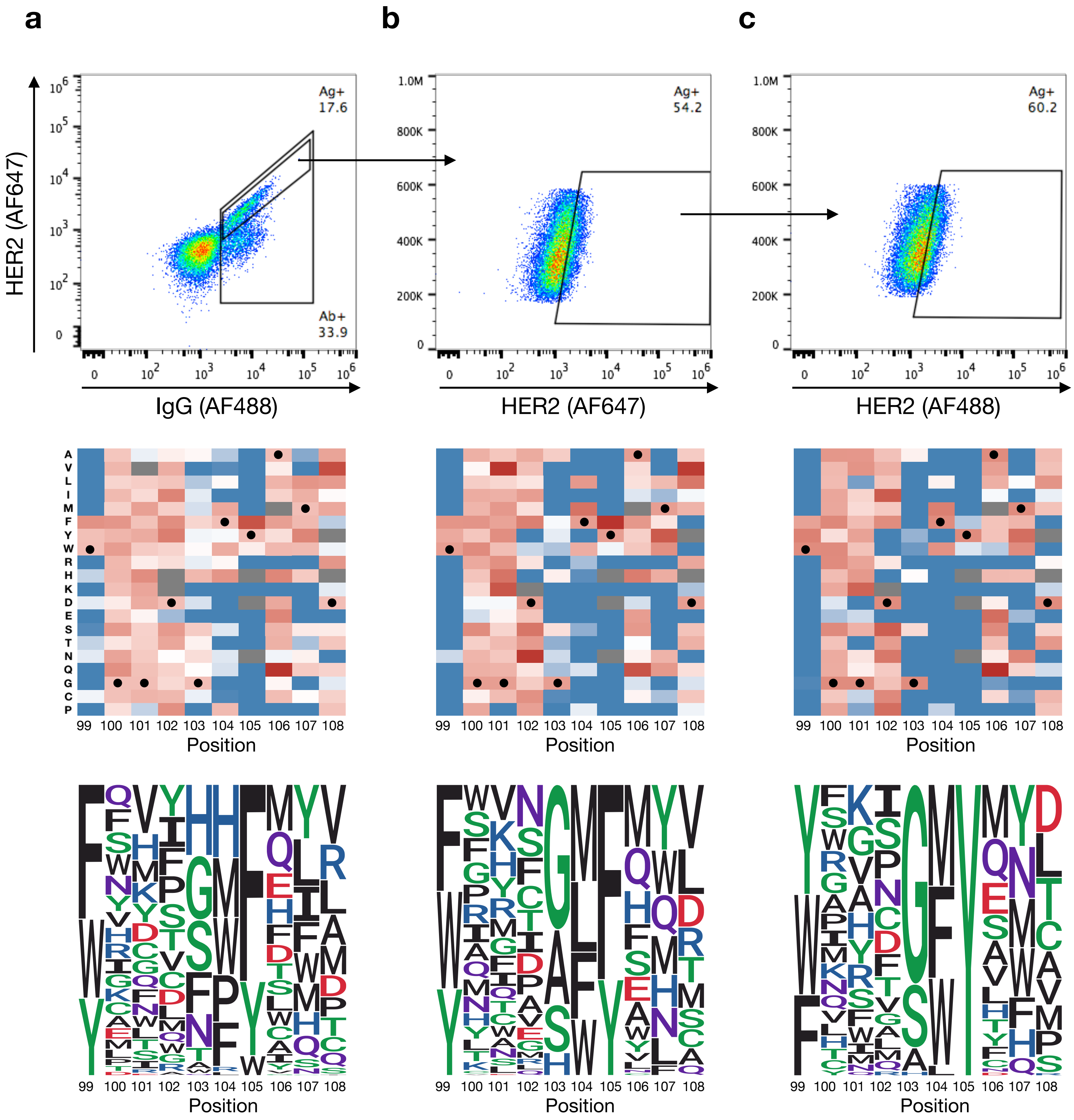


**Supplementary Figure 2: Trastuzumab CDRH3 deep mutational scanning**

Flow cytometry data for Trastuzumab (Herceptin) CDRH3 deep mutation scanning. (**a**) Flow cytometry plot, heatmap, and sequence logo plot following FACS for antibody expressing (Ab+) cells and antigen-specific (Ag+) cells. (**b**) Flow cytometry plot, heatmap, and sequence logo plot following a second round of enrichment for antigen-specific (Ag+2) cells; Decreased antigen concentration was used for flow cytometry labeling. (**c**) Flow cytometry plot, heatmap, and sequence logo plot following a third round of enrichment for antigen-specific (Ag+3) cells; Labeling for flow cytometry performed with antigen containing an alternatively conjugated fluorophore (Alexa Fluor 488). All enrichment ratios (ER) are calculated by dividing the frequency of a mutant found in the respective Ag+ population by the frequency of the mutant found in the Ab+ population.

| Sample | Description | Raw Read count (post-merge) | Aligned Reads | Unique CDRH3s |
|---|---|---|---|---|
| DMS-H3-Ab | Library of antibody expressing hybridomas following transfection with NNK-tiled mutagenesis ssODN donors | 1'060'062 | 992,772 (93.65%) | 2'786 |
| DMS-H3-Ag1 | Library of antigen specific antibodies expressed in hybridomas following transfection with NNK-tiled mutagenesis ssODN donors | 2'122'279 | 1,982,104 (93.4%) | 1'648 |
| DMS-H3-Ag2 | Library of antigen specific antibodies in hybridomas following two rounds of enrichment for antigen binding variants | 1'604'880 | 1,505,189 (93.79%) | 790 |
| DMS-H3-Ag3 | Library of antigen specific antibodies in hybridomas following three rounds of enrichment for antigen binding variants | 1'496'213 | 1,407,744 (94.09) | 663 |

**Supplementary Table 1: DMS deep sequencing statistics**

Deep sequencing results for sequence reads, alignment, and number of unique sequences detected for deep sequencing performed on the library (Ab+) and binding variants after 1, 2, 3 rounds of enrichment (Ag1, Ag2, Ag3).

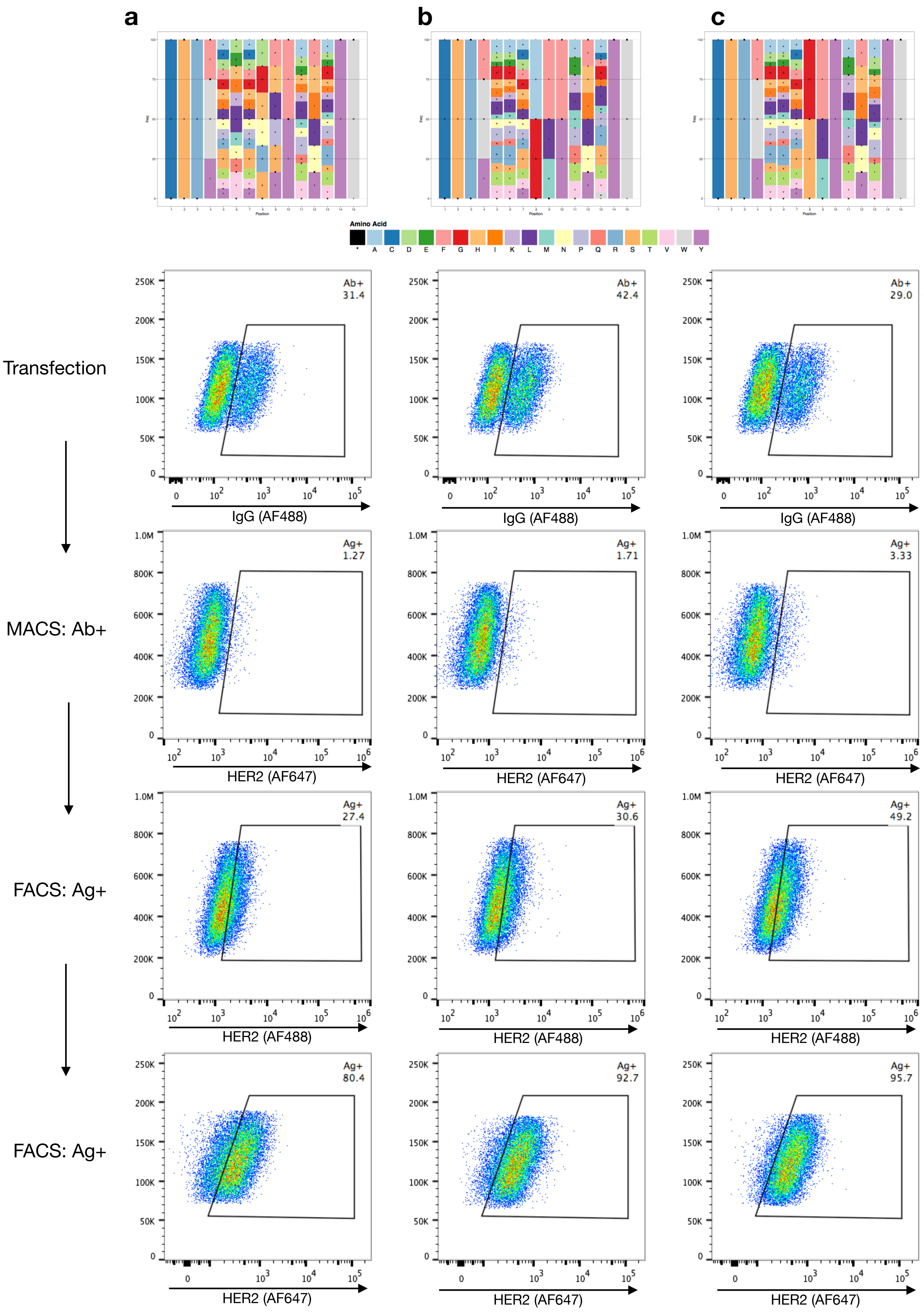


**Supplementary Figure 3: Generating antigen specific libraries in mammalian cells**

Workflow and flow cytometry data for generating antigen specific libraries in mammalian cells. First, libraries are generated by transfecting gRNA and ssODN donor templates containing rationally designed libraries. Second, antibody expressing cells (Ab+) are enriched by magnetic activated cell sorting (MACS). Ab+ cells can then undergo multiple rounds of enrichment for antigen-specific variants. (**a**) Libraries designed from DMS data following one round of antigen enrichment. (**b**) Libraries designed from DMS data following two rounds of antigen enrichment. (**c**) Libraries designed from DMS data following three rounds of antigen enrichment.

| Sample | Description | Raw Read count (post-merge) | Aligned Reads | Unique CDRH3s |
|---|---|---|---|---|
| CM-HER-H3-1-Ab | Antibody expressing hybridomas following transfection with ssODN donors for DMS-guided combinatorial mutagenesis library (DMS-Ag1) | 2'909'373 | 2,650,842 (91.11%) | 20'008 |
| CM-HER-H3-1-AgNeg | Variants sorted for negative antigen binding (non-binders) from the DMS-Ag1 DMS-guided combinatorial mutagenesis library. | 1'071'601 | 962,903 (89.86%) | 5'951 |
| CM-HER-H3-1-Ag1 | Variants sorted for positive antigen binding (binders) from the DMS-Ag1 DMS-guided combinatorial mutagenesis library. | 386'581 | 347,238 (89.82%) | 4'870 |
| CM-HER-H3-1-Ag2a | Variants following two rounds of FACS enrichment for positive antigen binding (binders) from the DMS-Ag1 DMS-guided combinatorial mutagenesis library; Second enrichment with AlexaFluor-647 conjugated HER2. | 644'907 | 587,013 (91.02%) | 3'345 |
| CM-HER-H3-1-Ag2b | Variants following two rounds of FACS enrichment for positive antigen binding (binders) from the DMS-Ag1 DMS-guided combinatorial mutagenesis library; Second enrichment with AlexaFluor-488 conjugated HER2. | 989'211 | 890,996 (90.07%) | 2'656 |
| CM-HER-H3-2-Ab | Antibody expressing hybridomas following transfection with ssODN donors for DMS-guided combinatorial mutagenesis library (DMS-Ag2) | 2'706'438 | 2,450,436 (90.54%) | 12'203 |
| CM-HER-H3-2-AgNeg | Variants sorted for negative antigen binding (non-binders) from the DMS-Ag2 DMS-guided combinatorial mutagenesis library. | 1'353'045 | 1,220,552 (90.21%) | 2'407 |
| CM-HER-H3-2-Ag1 | Variants sorted for positive antigen binding (binders) from the DMS-Ag2 DMS-guided combinatorial mutagenesis library. | 878'197 | 782,792 (89.14%) | 6'195 |
| CM-HER-H3-2-Ag2a | Variants following two rounds of FACS enrichment for positive antigen binding (binders) from the DMS-Ag2 DMS-guided combinatorial mutagenesis library; Second enrichment with AlexaFluor-647 conjugated HER2. | 819'196 | 722,903 (88.25%) | 4'192 |
| CM-HER-H3-2-Ag2b | Variants following two rounds of FACS enrichment for positive antigen binding (binders) from the DMS-Ag2 DMS-guided combinatorial mutagenesis library; Second enrichment with AlexaFluor-488 conjugated HER2. | 687'204 | 629,370 (91.58%) | 4'236 |
| CM-HER-H3-3-Ab | Antibody expressing hybridomas following transfection with ssODN donors for DMS-guided combinatorial mutagenesis library (DMS-Ag3) | 1'247'979 | 1,134,812 (90.93%) | 5'685 |
| CM-HER-H3-3-AgNeg | Variants sorted for negative antigen binding (non-binders) from the DMS-Ag3 DMS-guided combinatorial mutagenesis library. | 659'576 | 590,912 (89.59%) | 3'599 |
| CM-HER-H3-3-Ag1 | Variants sorted for positive antigen binding (binders) from the DMS-Ag3 DMS-guided combinatorial mutagenesis library. | 1'005'140 | 893,479 (88.89%) | 7'007 |
| CM-HER-H3-3-Ag2a | Variants following two rounds of FACS enrichment for positive antigen binding (binders) from the DMS-Ag3 DMS-guided combinatorial mutagenesis library; Second enrichment with AlexaFluor-647 conjugated HER2. | 728'831 | 646,857 (88.75%) | 5'502 |
| CM-HER-H3-3-Ag2b | Variants following two rounds of FACS enrichment for positive antigen binding (binders) from the DMS-Ag3 DMS-guided combinatorial mutagenesis library; Second enrichment with AlexaFluor-488 conjugated HER2. | 629'613 | 552,806 (87.80%) | 4'704 |

**Supplementary Table 2: DMS-combinatorial mutagenesis deep sequencing statistics**

Deep sequencing results for sequence reads, alignment, and number of unique sequences detected for deep sequencing performed on the library (Ab+), non-binding variants (AgNeg), and binding variants after 1 and 2 rounds of enrichment (Ag1, Ag2a, Ag2b).

**a** Model Performance: All data

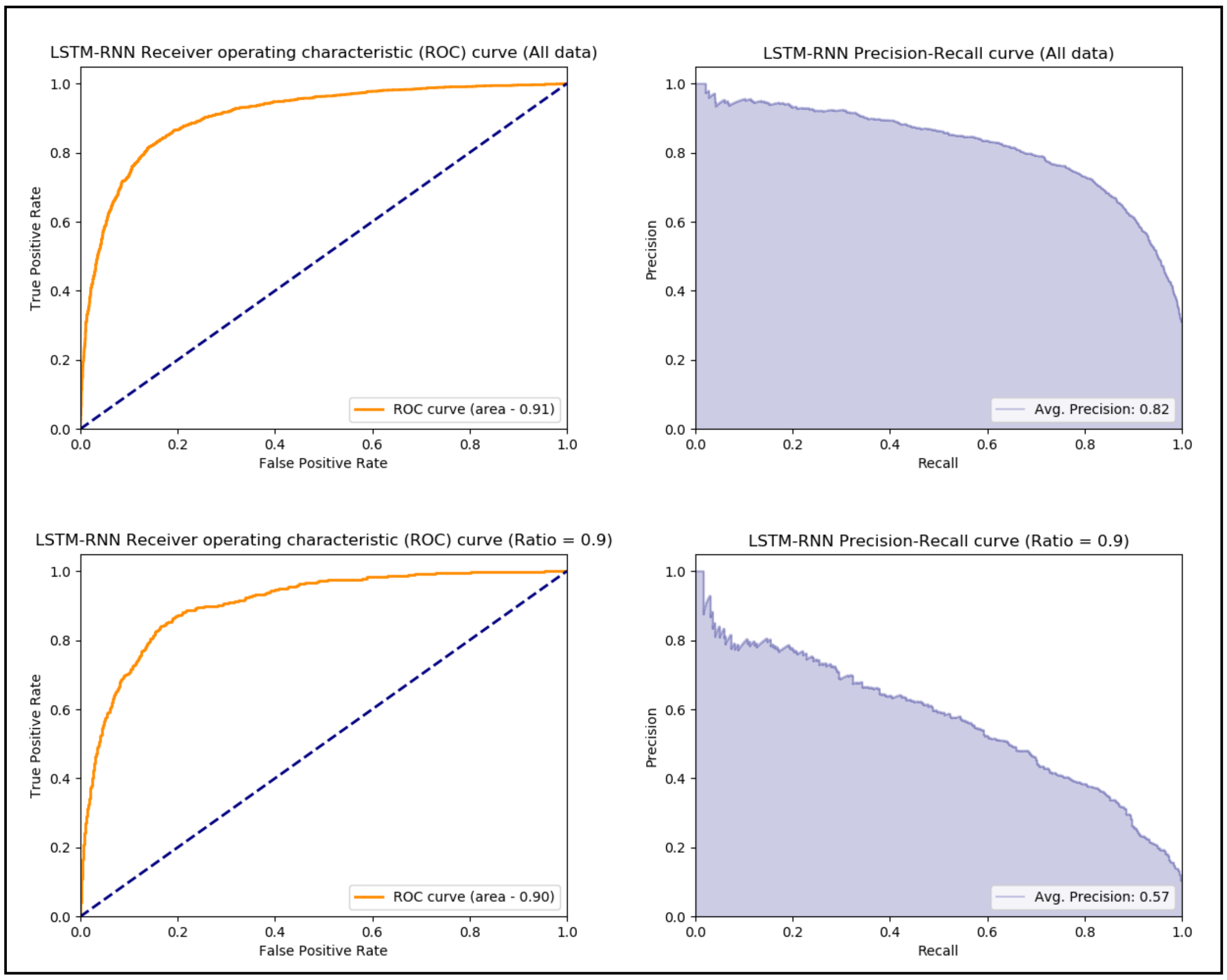


**b** Model Performance: All data

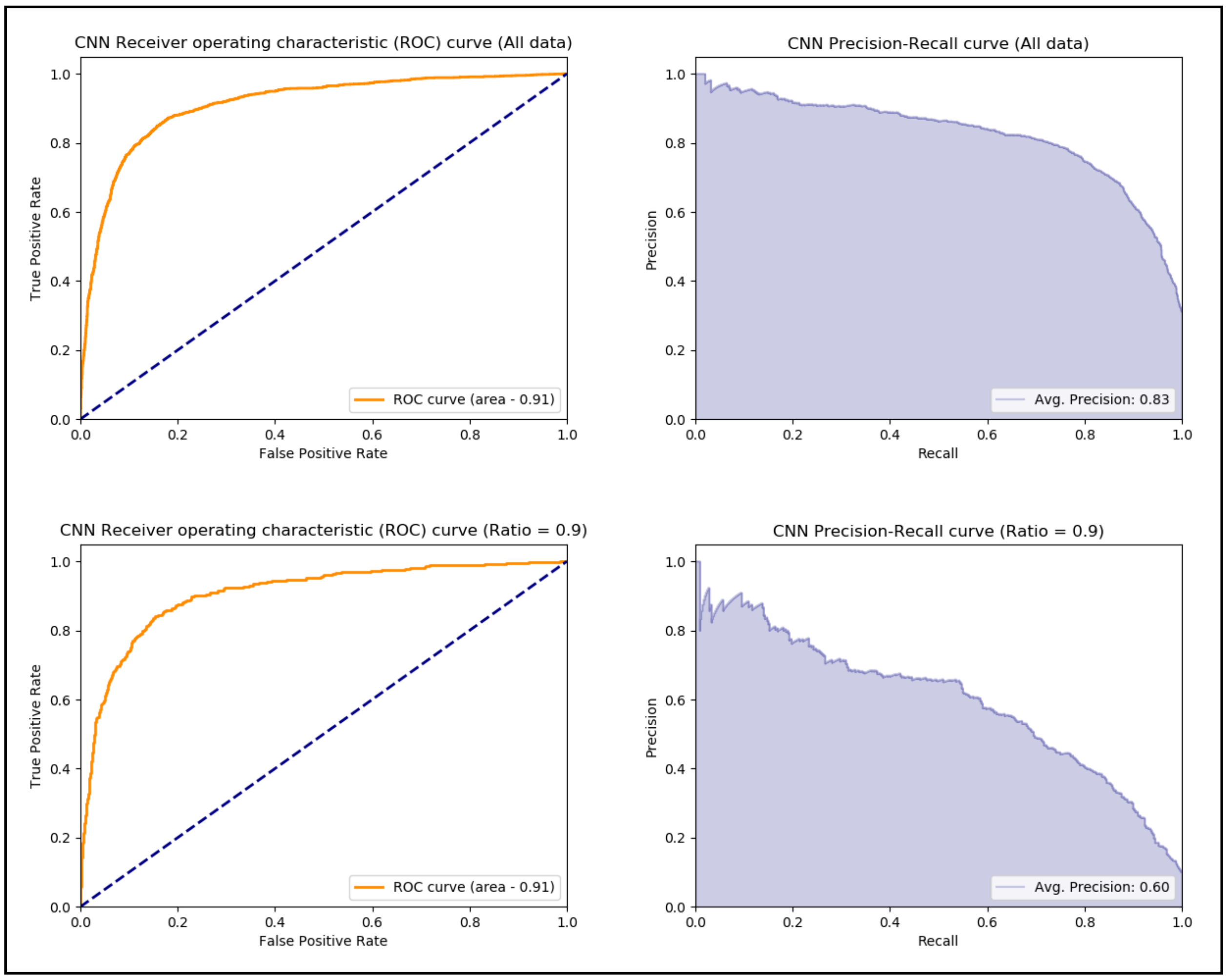


**Supplementary Figure 4: Model performance with all data**

**(a)** (top) LSTM-RNN ROC curve and PR curve for classification of withheld test data containing the same class split as seen in the training data set (top); In this instance, all available data resulted in a class split of approximately 30/70. (bottom) LSTM-RNN ROC curve and PR curve for classification of withheld test data containing a class split of approximately 10/90 to resemble physiologically expected frequencies based on flow cytometry data. **(b)** CNN ROC curves, PR curves as described above for the LSTM-RNN ROC and PR curves.

**a** Model Performance: 50/50 class split

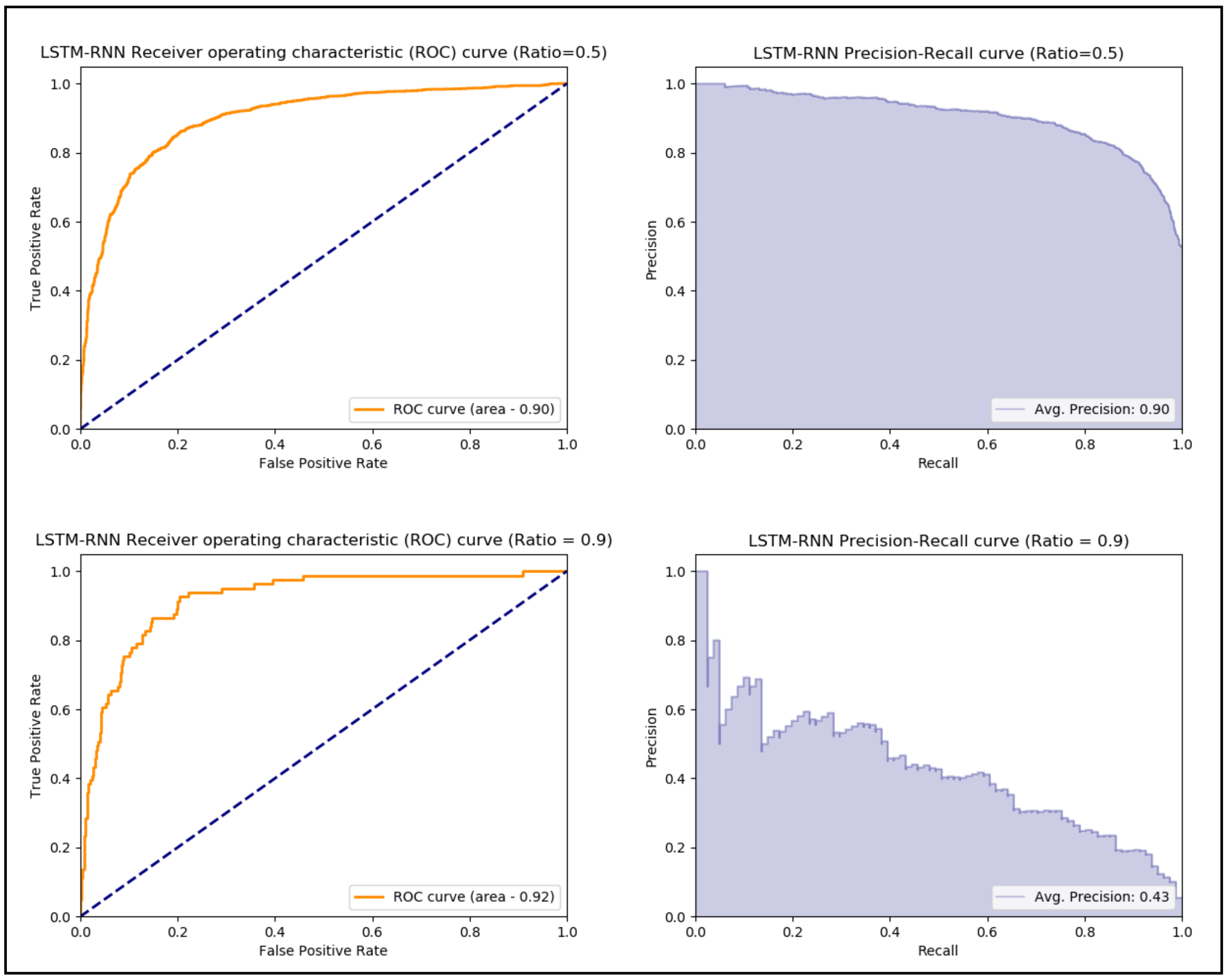


**b** Model Performance: 50/50 class split

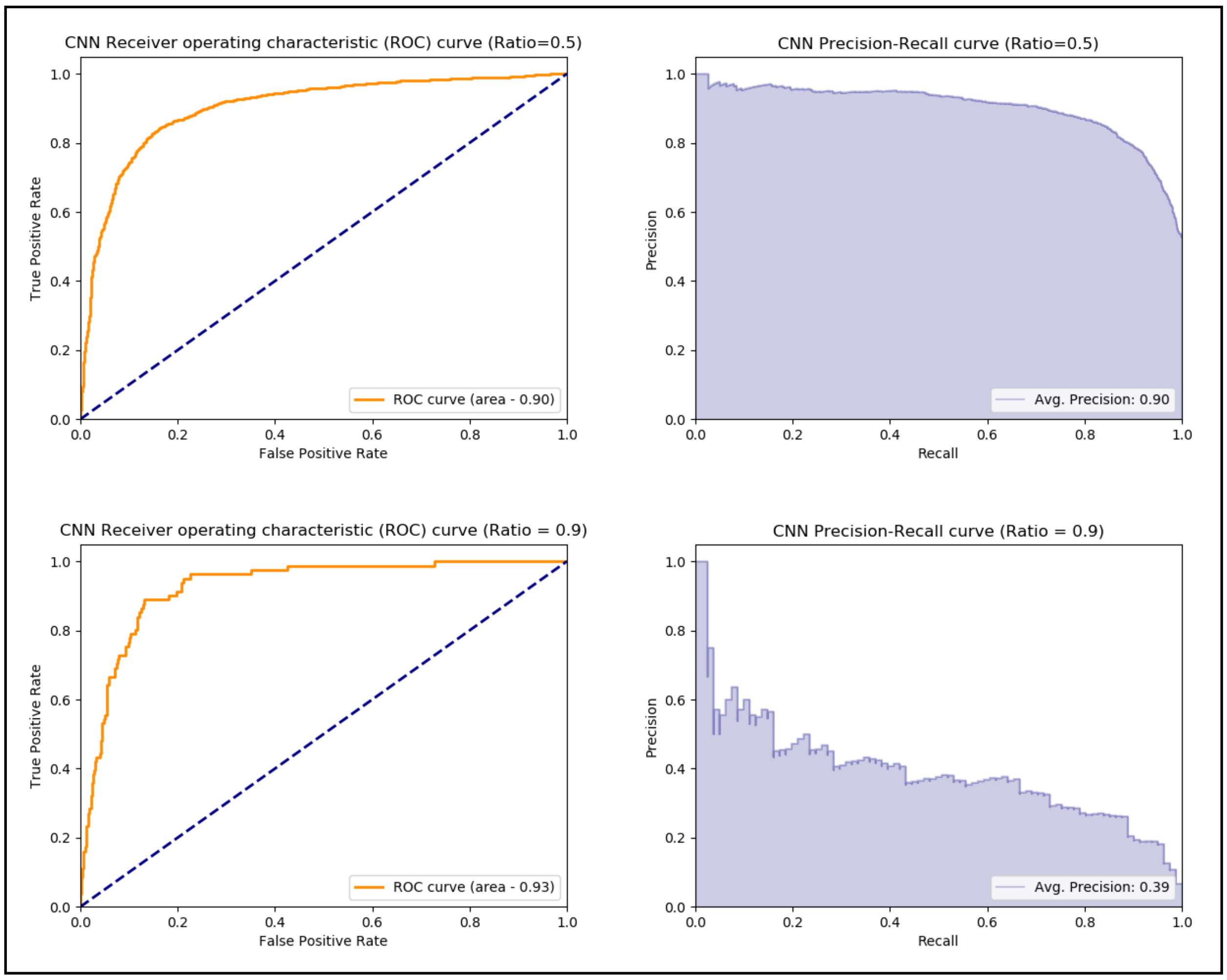


**Supplementary Figure 5: Model performance with a 50/50 class split data set**

**(a)** (top) LSTM-RNN ROC curve and PR curve for classification of withheld test data containing the same class split as seen in the training data set (top); In this instance, a class split of 50/50. (bottom) LSTM-RNN ROC curve and PR curve for classification of withheld test data containing a class split of approximately 10/90 to resemble physiologically expected frequencies based on flow cytometry data. **(b)** CNN ROC curves, PR curves as described above for the LSTM-RNN ROC and PR curves.

**a** Model Performance: 20/80 class split

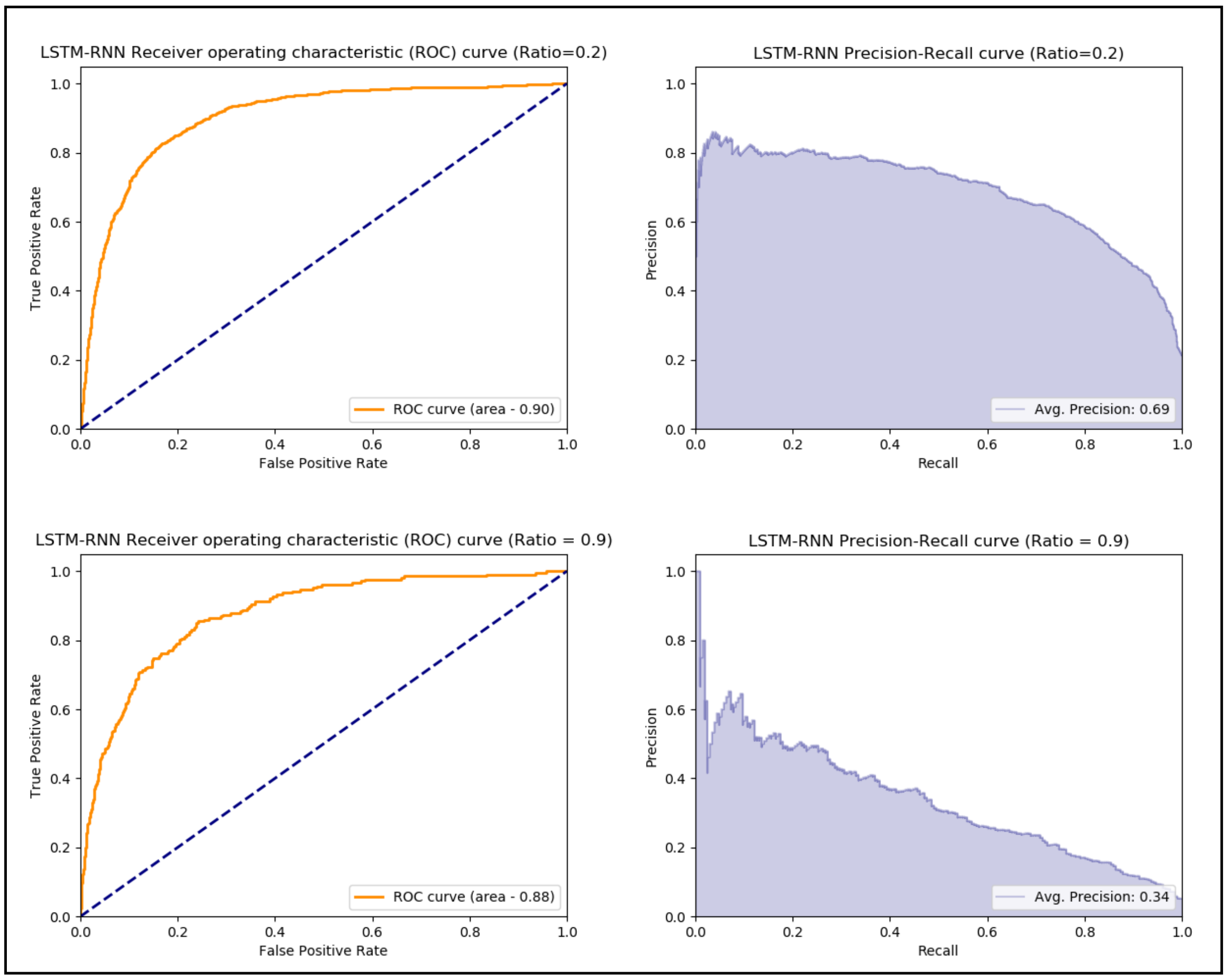


**b** Model Performance: 20/80 class split

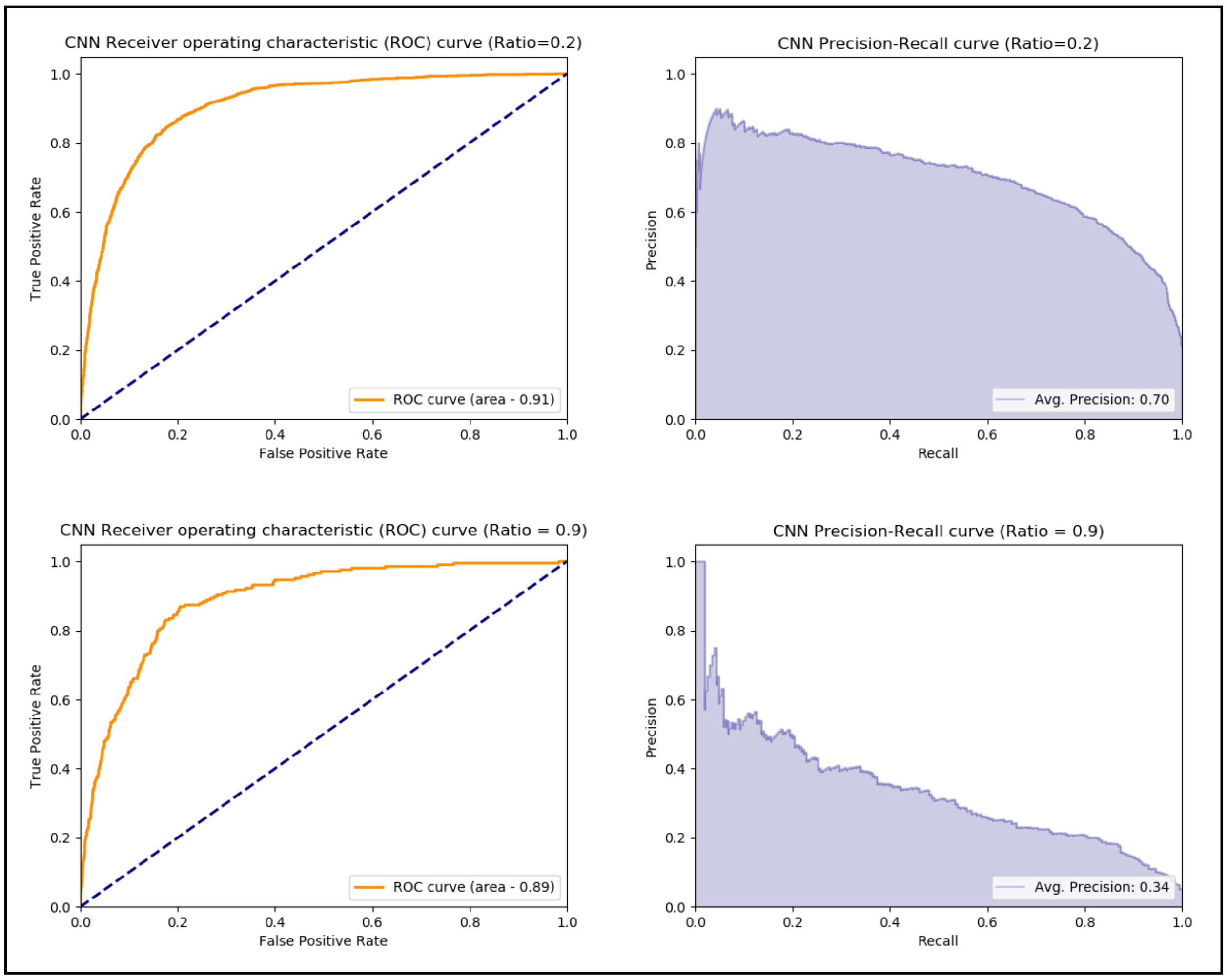


**Supplementary Figure 6: Model performance with a 20/80 class split data set**

**(a)** (top) LSTM-RNN ROC curve and PR curve for classification of withheld test data containing the same class split as seen in the training data set (top); In this instance, a class split of 20/80. (bottom) LSTM-RNN ROC curve and PR curve for classification of withheld test data containing a class split of approximately 10/90 to resemble physiologically expected frequencies based on flow cytometry data. **(b)** CNN ROC curves, PR curves as described above for the LSTM-RNN ROC and PR curves.

**a** Model Performance: 10/90 class split

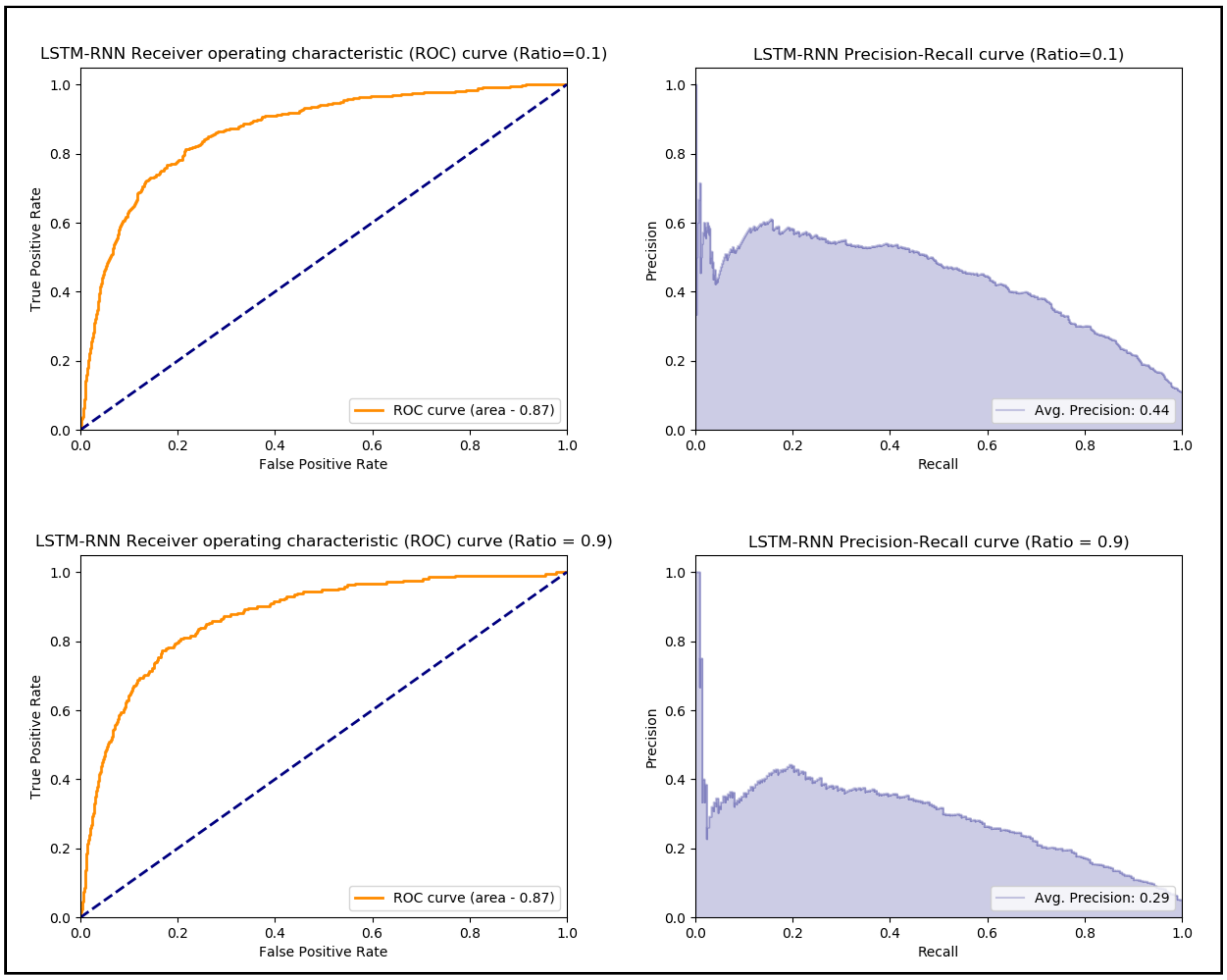


**b** Model Performance: 10/90 class split

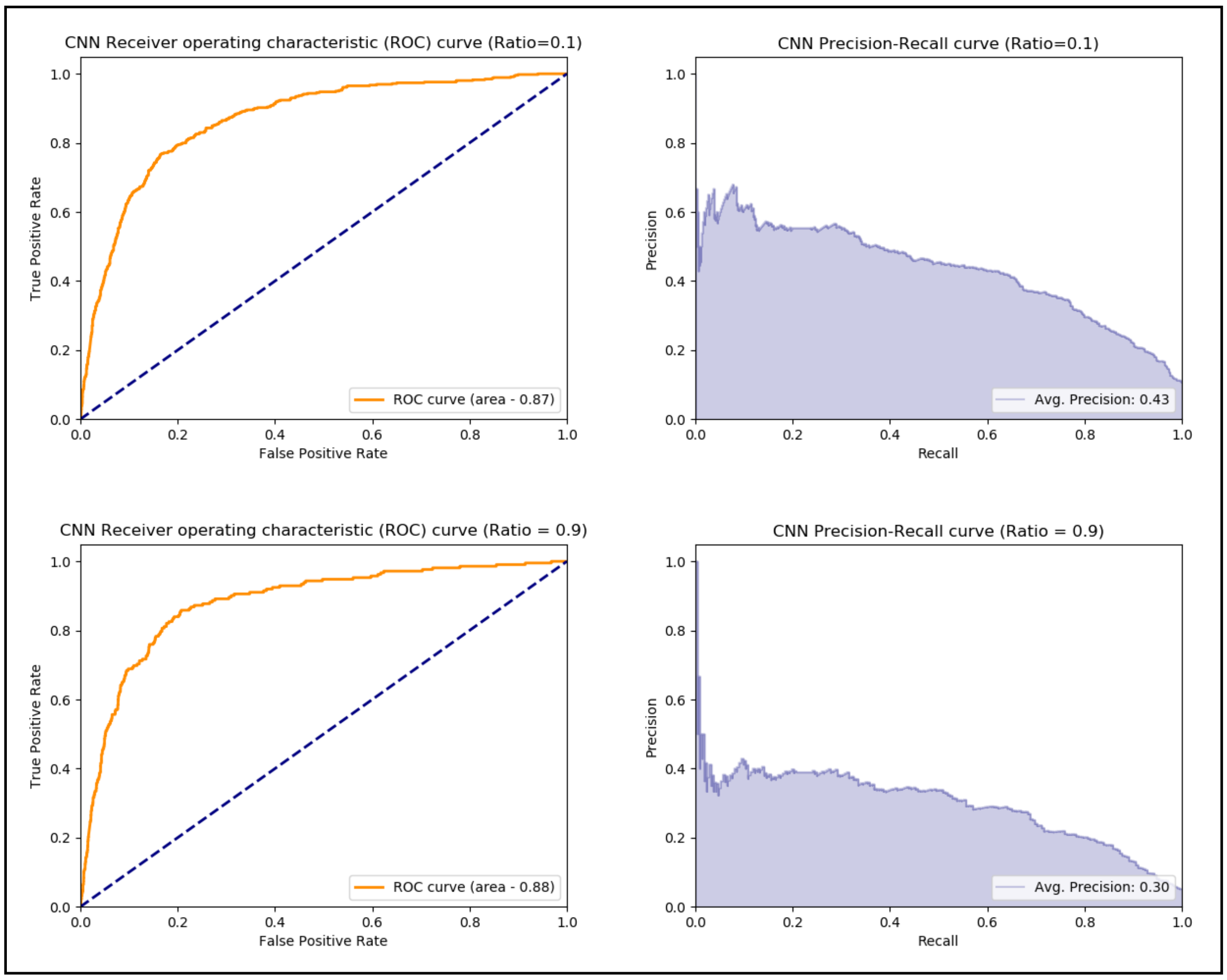


**Supplementary Figure 7: Model performance with a 10/90 class split data set**

**(a)** (top) LSTM-RNN ROC curve and PR curve for classification of withheld test data containing the same class split as seen in the training data set (top); In this instance, a class split of 10/90. (bottom) LSTM-RNN ROC curve and PR curve for classification of withheld test data containing a class split of approximately 10/90 to resemble physiologically expected frequencies based on flow cytometry data. **(b)** CNN ROC curves, PR curves as described above for the LSTM-RNN ROC and PR curves.

**a** Model Performance: Random shuffled labels

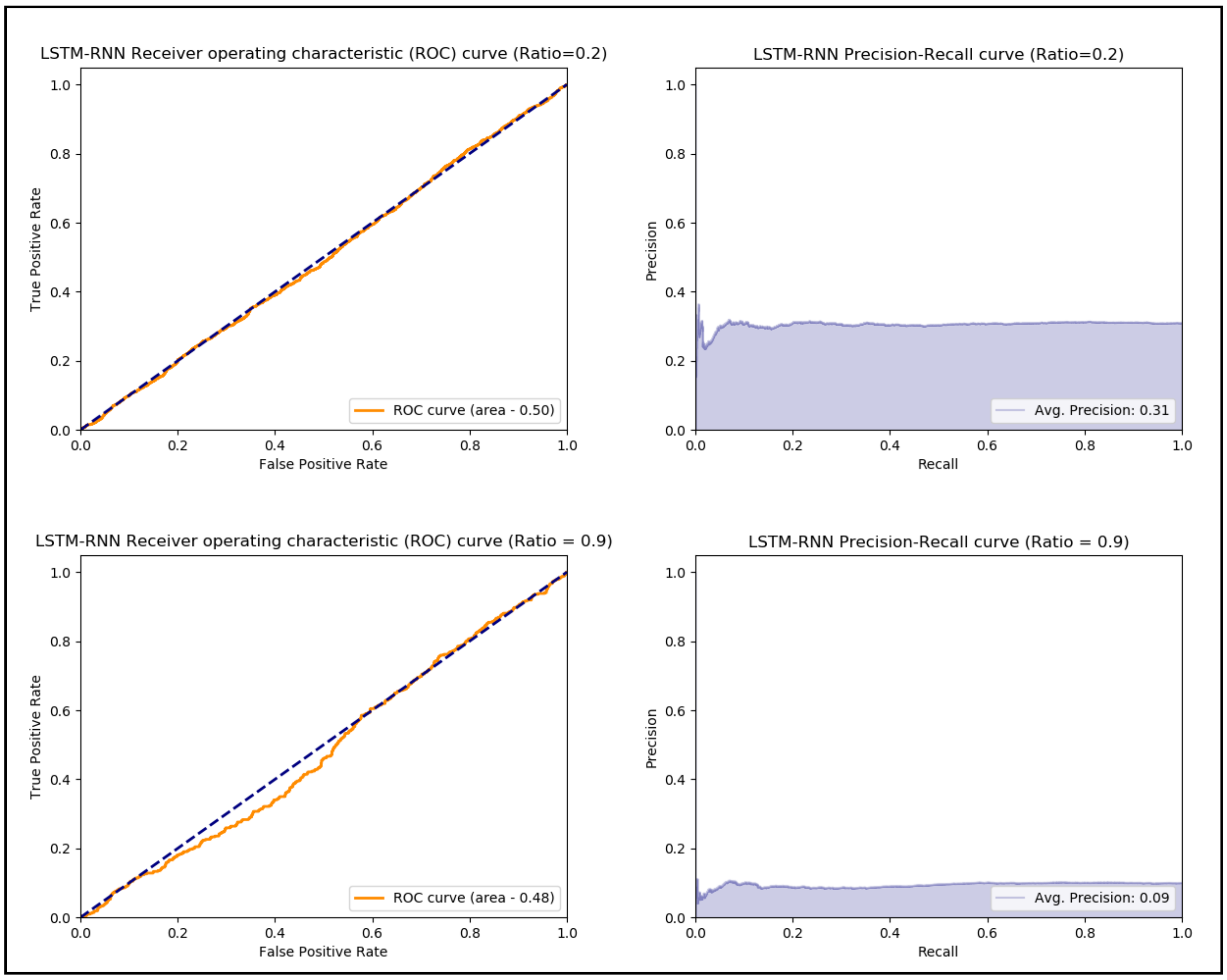


**b** Model Performance: Random shuffled labels

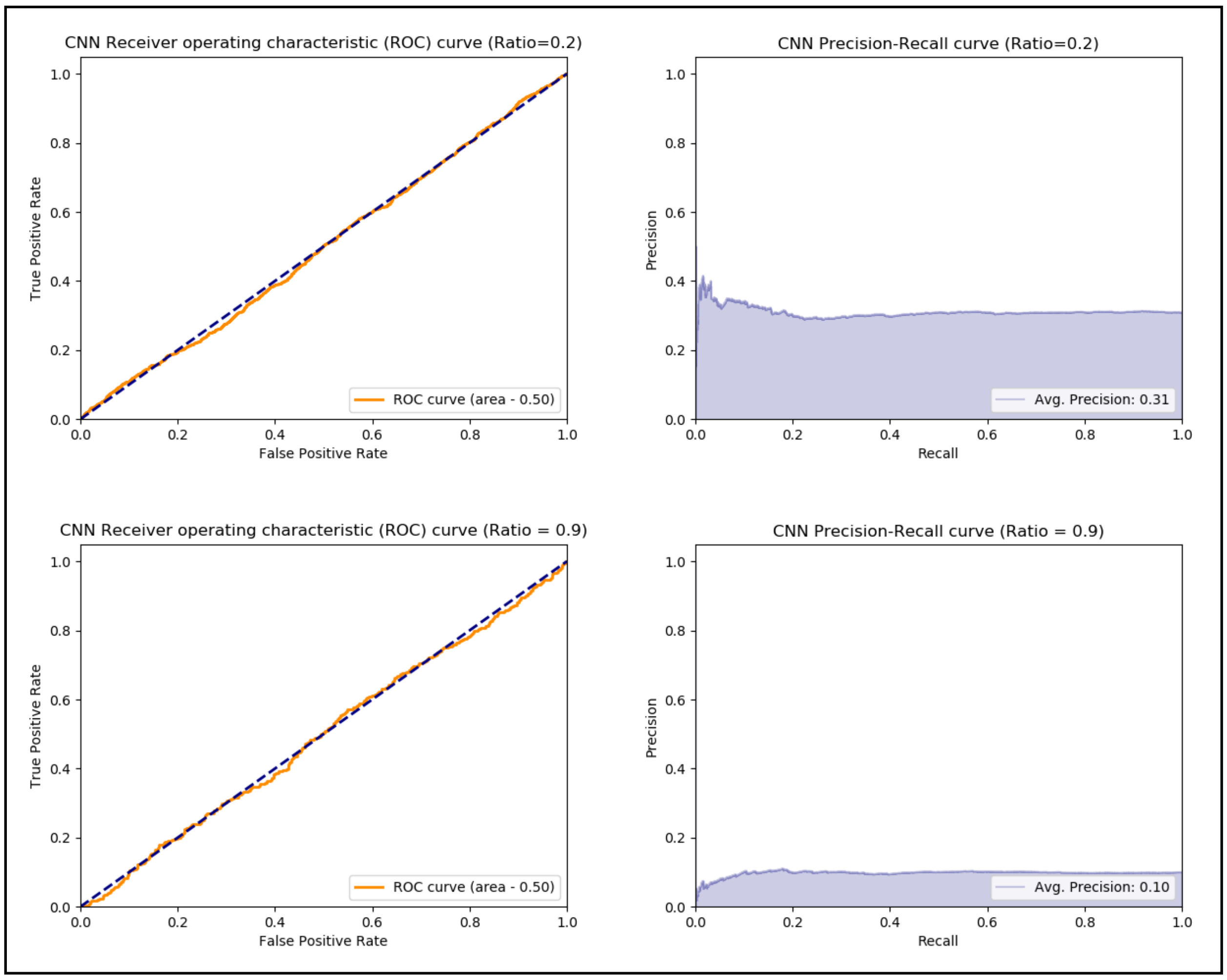


**Supplementary Figure 8: Model performance on randomly shuffled labels**

Randomly shuffling binding/non-binding class labels produces neural networks incapable of learning or identifying any associated patterns. **(a)** (top) LSTM-RNN ROC curve and PR curve for classification of withheld test data containing the same class split as seen in the training data set (top); In this instance, all available data resulted in a class split of approximately 30/70. (bottom) LSTM-RNN ROC curve and PR curve for classification of withheld test data containing a class split of approximately 10/90 to resemble physiologically expected frequencies based on flow cytometry data. **(b)** CNN ROC curves, PR curves as described above for the LSTM-RNN ROC and PR curves.

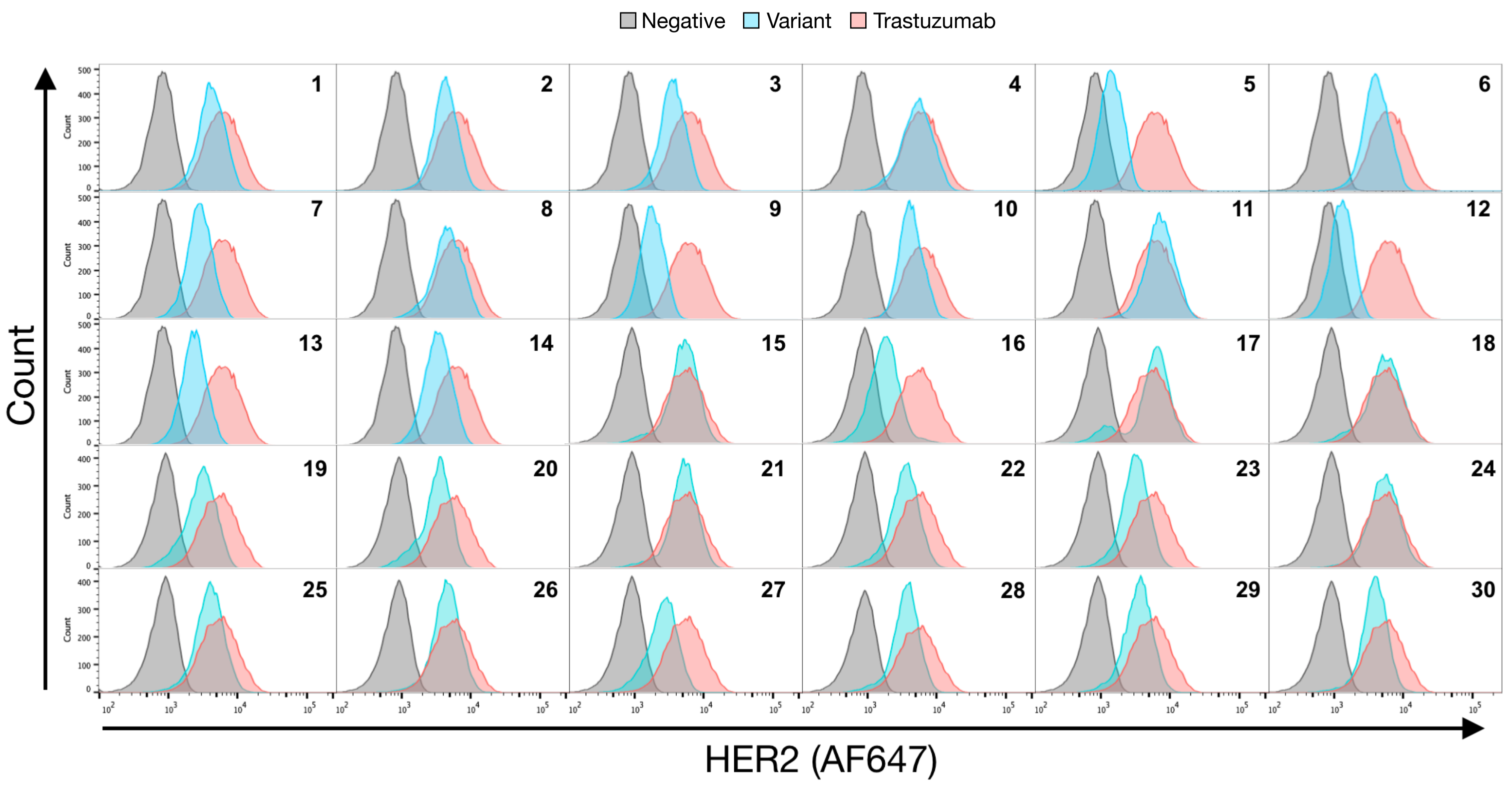


**Supplementary Figure 9: Flow cytometry profiles for the 30 randomly selected variants**

Flow cytometry profiles for the individual sequence variants confirms binding activity for 100% (30/30) of the sequences.

**a**

| Model / Class split | ROC curve AUC (Train Split) | Average PR (Train Split) | ROC curve AUC (10/90 Split) | Average PR (10/90 Split) |
|---|---|---|---|---|
| RNN All data | 0.91 | 0.82 | 0.90 | 0.57 |
| CNN All data | 0.91 | 0.83 | 0.91 | 0.60 |
| RNN 50/50 | 0.90 | 0.90 | 0.92 | 0.43 |
| CNN 50/50 | 0.90 | 0.90 | 0.93 | 0.39 |
| RNN 20/80 | 0.90 | 0.69 | 0.88 | 0.34 |
| CNN 20/80 | 0.91 | 0.70 | 0.89 | 0.34 |
| RNN 10/90 | 0.87 | 0.44 | 0.87 | 0.29 |
| CNN 10/90 | 0.87 | 0.43 | 0.88 | 0.30 |

**b**

| Model / Class split | Predicted Binders |
|---|---|
| RNN All data | 17'489 |
| CNN All data | 17'930 |
| RNN 50/50 | 26'480 |
| CNN 50/50 | 24'156 |
| RNN 20/80 | 13'893 |
| CNN 20/80 | 11'336 |
| RNN 10/90 | 4'296 |
| CNN 10/90 | 8'041 |

**Supplementary Table 3: Model metric summary and prediction estimates**

**(a)** Summary table of the performance characteristics for the LSTM-RNN and CNN trained and tested on data sets with various ratios or class splits of binding/non-binding sequences. Model training was performed on 70% of the data and testing was performed by withholding the remaining 30% and then comparing the model's classification of test sequences with the known classification. The withheld sequences were further split into two test sets; One test set contained the same class split as the training data (Train Split) and the other contained a class split of approximately 10/90 to resemble physiologically expected frequencies (10/90 Split). **(b)** 100,000 sequences were generated by randomly sampling amino acids from the expected usage based on the combinatorial mutagenesis scheme and classified by the trained LSTM-RNN and CNN. The class split of the training data substantially impacts the model's classification of binders/non-binders.

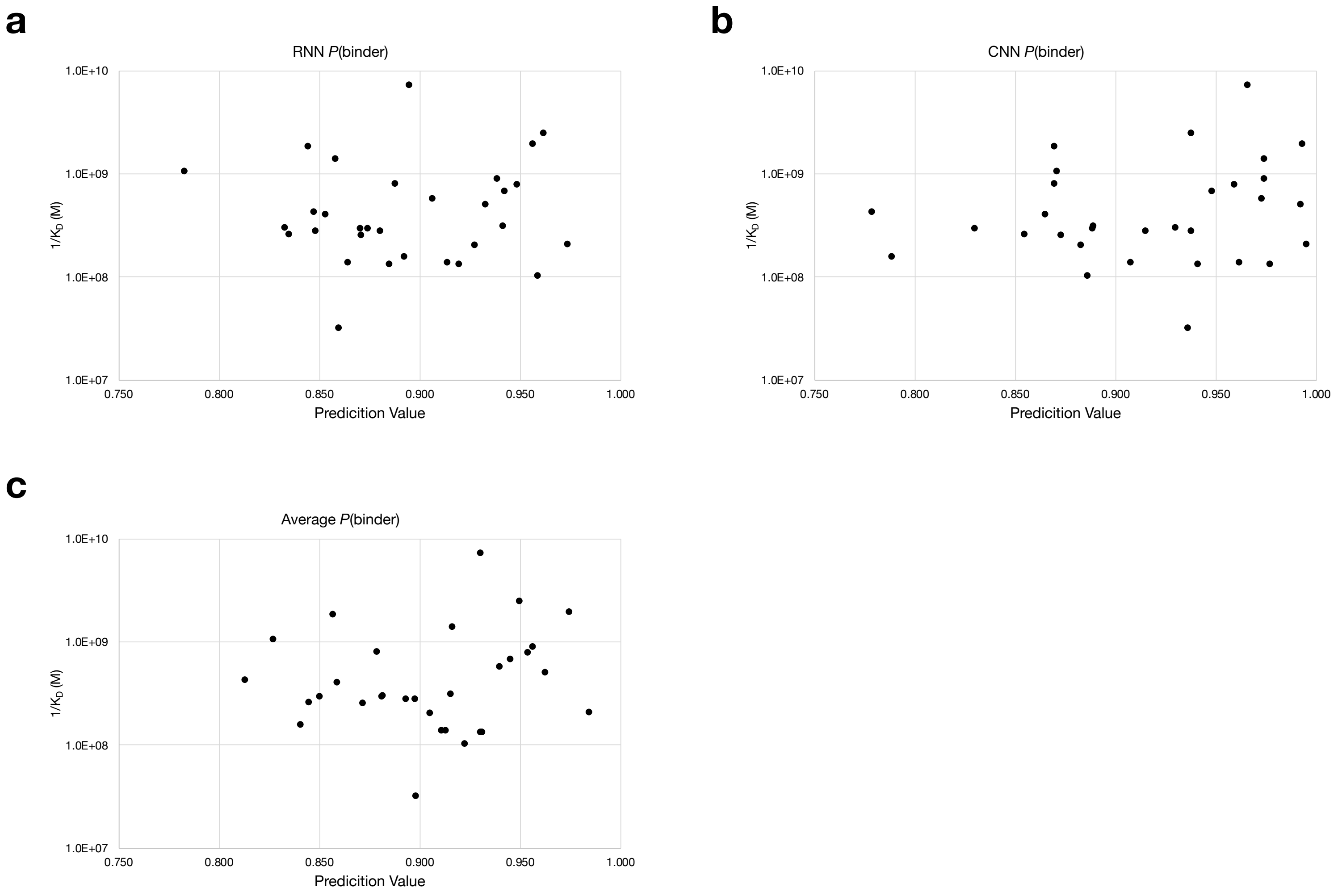


**Supplementary Figure 10: Affinity values plotted against their binding prediction values**

**(a)** Affinity values plotted against the LSTM-RNN prediction values. (**b)** Affinity values plotted against the CNN prediction values. **(c)** Affinity values plotted against the LSTM-RNN and CNN averaged prediction values. No observable trends appear to be present, especially in the context of the LSTM-RNN prediction values, however, for the CNN and LSTM-RNN/CNN averaged prediction values, higher affinity variants tend to have higher prediction values.

| LD | % of sequences in largest cluster | Negative predictions in largest cluster | Positive predictions in largest cluster | % positive predictions in largest cluster |
|---|---|---|---|---|
| 1 | 0.03% | 0 | 3 | 100% |
| 2 | 30.0% | 259 | 2743 | 91.2% |
| 3 | 99.7% | 4971 | 5000 | 50.1% |
| 4 | 100% | 5000 | 5000 | 50.0% |
| 5 | 100% | 5000 | 5000 | 50.0% |
| 6 | 100% | 5000 | 5000 | 50.0% |

**Supplementary Table 4: Network analysis of predicted binders and non-binders**

5,000 predicted binding and 5,000 predicted non-binding sequences were randomly selected and networks were generated by clustering sequences at Levenshtein distances (*LD*) up to 6. For the network at *LD* ≤ 2 the majority of sequences (54%) do not form clusters or form a cluster with a single other sequence.

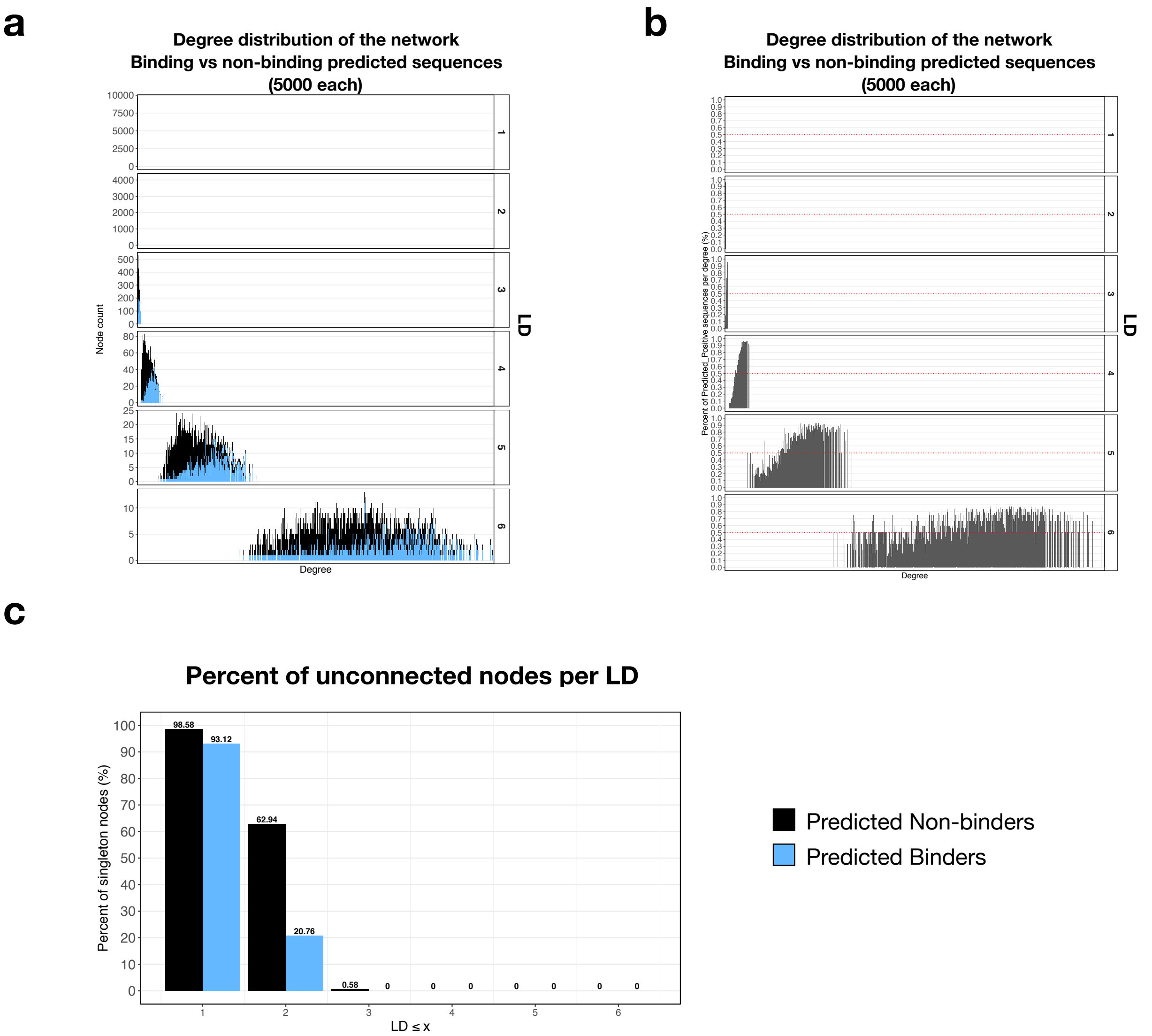


**Supplementary Figure 11: Network analysis of predicted binders and non-binders**

**(a)** For networks *LD* 1-6: The counts of positive and negative sequences that have a given degree. Negative predicted sequences are overrepresented in the lower degree bin meaning each node has less neighbors at any given distance than positive predicted sequences. **(b)** For networks LD 1-6: Percentage of predicted positive sequences for each degree count. The ratio of predicted positive to predicted negative increases as degrees are increased, showing again that predicted positive sequences are more connected. **(c)** The set of predicted negative sequences has a larger percentage of 'more different' sequences that are edit distance 1, 2, or 3 away from any other of the 10,000 sequences.

**a**

| Target/ Antigen | Working conc. | Incubation volume | Fluorophore | Product ID |
|---|---|---|---|---|
| Human IgG (H+L) | 1.5 ug/mL | 100 µl | AlexaFluor® 488 | 109-545-088 (Jackson ImmunoResearch) |
| ErbB2 (HER2) Round 1 | 2.9 µg/ml (40 nM) | 100 µl | AlexaFluor® 647 | SRP6405-50UG (Sigma-Aldrich) |
| ErbB2 (HER2) Round 2 | 1.4 µg/ml (20 nM) | 100 µl | AlexaFluor® 647 | SRP6405-50UG (Sigma-Aldrich) |
| ErbB2 (HER2) Round 3 | 1.4 µg/ml (20 nM) | 100 µl | AlexaFluor® 488 | SRP6405-50UG (Sigma-Aldrich) |

**b**

| Target/ Antigen | Working conc. | Incubation volume | Fluorophore | Product ID |
|---|---|---|---|---|
| Human IgG (H+L) | 1.5 ug/mL | 100 µl | AlexaFluor® 488 | 109-545-088 (Jackson ImmunoResearch) |
| ErbB2 (HER2) Round 1 | 2.9 µg/ml (40 nM) | 100 µl | AlexaFluor® 647 | SRP6405-50UG (Sigma-Aldrich) |
| ErbB2 (HER2) Round 2a | 1.4 µg/ml (20 nM) | 100 µl | AlexaFluor® 488 | SRP6405-50UG (Sigma-Aldrich) |
| ErbB2 (HER2) Round 2b | 1.4 µg/ml (20 nM) | 100 µl | AlexaFluor® 647 | SRP6405-50UG (Sigma-Aldrich) |

**Supplementary Table 5: Flow cytometry labeling conditions**

**(a)** Flow cytometry labeling conditions for deep mutational scanning studies. **(b)** Flow cytometry labeling conditions for DMS-guided combinatorial mutagenesis libraries.

| Description | Sequence |
|---|---|
| Trastuzumab variant gRNA target sequence | GCCCCAATAATCGAACCCAGAGG |
| VH forward primer for NGS preparation via primer extension (PCR1) | CCCTCCTTTAATTCCCCTTTTTGTCTAAAGGTATCCTGTCA |
| VH reverse primer for NGS preparation via primer extension (PCR1) | GAGGAGAGAGAGAGAGGACCTGGAGAGGCCATTC |
| TruSeq Universal Adapter forward primer for NGS preparation via primer extension (PCR2) | UNIVERSAL ADAPTER - NNNN CCCTCCTTTAATTCCC |
| TruSeq Index Adapter reverse primer for NGS preparation via primer extension (PCR2) | ADAPTER INDEX X - NNNN GAGGAGAGAGAGAGAG |

**Supplementary Table 6: gRNA target and primer sequences**

A summary table providing the nucleotide sequences of all relevant primers and gRNA target sites referenced in this study.

| Description | Sequence |
|---|---|
| 5' homology arm-<br>CDRH3 sequence<br>-3' homology arm | CTGAGAGCCGAGGACACGGCTGTGTATTACTGTTCGCGA-<br>TGGGGAGGTGATGGATTTTATGCAATGGAT<br>-TACTGGGGCCAGGGAACCCTGGTCACCGTCTCCTCAGGTAAG |
| Example sequence for single-site mutagenesis. NNK sequence is tiled along the entire CDRH sequence | CTGAGAGCCGAGGACACGGCTGTGTATTACTGTTCGCGA-<br>**NNK** GGA GGT GAT GGA TTT TAT GCA ATG GAT<br>-TACTGGGGCCAGGGAACCCTGGTCACCGTCTCCTCAGGTAAG |
| Library designed off enrichment ratios from DMS-H3-Ag1; Combinatorial mutagenesis ssODN donor (1) | CAAATGAACAGCCTGAGAGCCGAGGACACGGCTGTGTATTACTGTAGCAGG-<br>**TWC NNC VWS NNC VRC YHC TWC VHS HWC VNS**<br>-TATTGGGGCCAGGGAACCCTGGTCACCGTCTCCTCAGGTAAGAATG |
| Library designed off enrichment ratios from DMS-H3-Ag1; Combinatorial mutagenesis ssODN donor (2) | CAAATGAACAGCCTGAGAGCCGAGGACACGGCTGTGTATTACTGTAGCAGG-<br>**TGG NNC VWS NNC VRC YHC TWC VHS HWC VNS**<br>-TATTGGGGCCAGGGAACCCTGGTCACCGTCTCCTCAGGTAAGAATG |
| Library designed off enrichment ratios from DMS-H3-Ag2; Combinatorial mutagenesis ssODN donor (1) | CAAATGAACAGCCTGAGAGCCGAGGACACGGCTGTGTATTACTGTAGCAGG-<br>**TWC VNB VNS NNC GST TTC TWC VHG HWC NBS**<br>-TATTGGGGCCAGGGAACCCTGGTCACCGTCTCCTCAGGTAAGAATG |
| Library designed off enrichment ratios from DMS-H3-Ag2; Combinatorial mutagenesis ssODN donor (2) | CAAATGAACAGCCTGAGAGCCGAGGACACGGCTGTGTATTACTGTAGCAGG-<br>**TGG VNB VNS NNC GST TTC TWC VHG HWC NBS**<br>-TATTGGGGCCAGGGAACCCTGGTCACCGTCTCCTCAGGTAAGAATG |
| Library designed off enrichment ratios from DMS-H3-Ag2; Combinatorial mutagenesis ssODN donor (3) | CAAATGAACAGCCTGAGAGCCGAGGACACGGCTGTGTATTACTGTAGCAGG-<br>**TWC VNB VNS NNC GST MTG TWC VHG HWC NBS**<br>-TATTGGGGCCAGGGAACCCTGGTCACCGTCTCCTCAGGTAAGAATG |
| Library designed off enrichment ratios from DMS-H3-Ag2; Combinatorial mutagenesis ssODN donor (4) | CAAATGAACAGCCTGAGAGCCGAGGACACGGCTGTGTATTACTGTAGCAGG-<br>**TGG VNB VNS NNC GST MTG TWC VHG HWC NBS**<br>-TATTGGGGCCAGGGAACCCTGGTCACCGTCTCCTCAGGTAAGAATG |
| Library designed off enrichment ratios from DMS-H3-Ag3; Combinatorial mutagenesis ssODN donor (1) | CAAATGAACAGCCTGAGAGCCGAGGACACGGCTGTGTATTACTGTAGCAGG-<br>**TWC VNB VNS NNC RGY TTC TAC VHG HWC VHB**<br>-TATTGGGGCCAGGGAACCCTGGTCACCGTCTCCTCAGGTAAGAATG |
| Library designed off enrichment ratios from DMS-H3-Ag3; Combinatorial mutagenesis ssODN donor (2) | CAAATGAACAGCCTGAGAGCCGAGGACACGGCTGTGTATTACTGTAGCAGG-<br>**TGG VNB VNS NNC RGY TTC TAC VHG HWC VHB**<br>-TATTGGGGCCAGGGAACCCTGGTCACCGTCTCCTCAGGTAAGAATG |
| Library designed off enrichment ratios from DMS-H3-Ag3; Combinatorial mutagenesis ssODN donor (3) | CAAATGAACAGCCTGAGAGCCGAGGACACGGCTGTGTATTACTGTAGCAGG-<br>**TWC VNB VNS NNC RGY MTG TAC VHG HWC VHB**<br>-TATTGGGGCCAGGGAACCCTGGTCACCGTCTCCTCAGGTAAGAATG |
| Library designed off enrichment ratios from DMS-H3-Ag3; Combinatorial mutagenesis ssODN donor (4) | CAAATGAACAGCCTGAGAGCCGAGGACACGGCTGTGTATTACTGTAGCAGG-<br>**TGG VNB VNS NNC RGY MTG TAC VHG HWC VHB**<br>-TATTGGGGCCAGGGAACCCTGGTCACCGTCTCCTCAGGTAAGAATG |

**Supplementary Table 7: ssODN donor sequences**

A summary table providing the nucleotide sequences of all relevant ssODN donors referenced in this study.
